# *Arabidopsis* cell wall composition determines disease resistance specificity and fitness

**DOI:** 10.1101/2020.05.21.105650

**Authors:** Antonio Molina, Eva Miedes, Laura Bacete, Tinguaro Rodríguez, Hugo Mélida, Nicolas Denancé, Andrea Sánchez-Vallet, Marie-Pierre Rivière, Gemma López, Amandine Freydier, Xavier Barlet, Sivakumar Pattathil, Michael Hahn, Deborah Goffner

## Abstract

Plant cell walls are complex structures subject to dynamic remodeling in response to developmental and environmental cues, and play essential functions in disease resistance responses. We tested the specific contribution of plant cell walls to immunity by determining the susceptibility of a set of *Arabidopsis* cell wall mutants (*cwm*) to pathogens with different parasitic styles: a vascular bacterium, a necrotrophic fungus and a biotrophic oomycete. Remarkably, most *cwm* mutants tested (31/38; 81.6%) showed alterations in their resistance responses to at least one of these pathogens, in comparison to wild-type plants, illustrating the relevance of wall composition in determining disease resistance phenotypes. We found that the enhanced resistance of *cwm* plants to the necrotrophic and vascular pathogens negatively impacted on *cwm* fitness traits, like biomass and seed yield. Enhanced resistance of *cwm* plants is not only mediated by canonical immune pathways, like those modulated by phytohormones or Microbe-Associated Molecular Patterns, which are not de-regulated in all *cwm* tested. Pectin-enriched wall fractions isolated from *cwm* plants triggered immune responses in other plants, suggesting that wall-mediated defensive pathways might contribute to *cwm* resistance. Cell walls of *cwm* plants show a high diversity of composition alterations as revealed by glycome profiling that detect specific wall carbohydrate moieties. Mathematical analysis of glycome profiling data identified correlations between the amounts of specific wall carbohydrate moieties and disease resistance phenotypes of *cwm* plants. These data support the relevant and specific function of plant wall composition in plant immune response modulation and in balancing disease resistance/development trade-offs.

---

Plants are under continuous pathogen threats that might compromise their survival and reproduction. To cope with these threats, plants have evolved a plethora of resistance mechanisms, which are either constitutively expressed or induced after pathogen attack (1–4). One common resistance mechanism to all plant cells is the presence of a cell wall that shields plants from pathogen invasion. The cell wall acts first as a passive barrier that pathogens have to hydrolyze, by secreting cell wall degrading enzymes, for infection progression, but also functions as a reservoir of antimicrobial compounds (5–7). Plant cell walls are also a source of carbohydrate moieties that are released during wall degradation and could act as Damage-Associated Molecular Patterns (DAMPs) triggering plant immune responses upon their perception by plants pattern recognition receptors (PRRs; 6–12). Plant walls are complex and dynamic structures that consist of a primary wall composed of carbohydrate-based polymers –cellulose, pectic polysaccharides (homogalacturonan, rhamnogalacturonan (RGI) and RGII), hemicelluloses (xyloglucan and xylans) and minor polysaccharides– and of structural glycoproteins (13). In addition, some plant cells to reinforce their structure deposit a secondary wall, that is mainly composed of cellulose, hemicelluloses (mostly xylans) and lignin (14, 15). The biosynthesis, transport, deposition, remodeling and turnover of cell walls, along with the regulation of these processes involve approximately 10% of genes encoded in plants genomes (16, 17).

Modifications of cell wall composition and structure occur during plant development, but also upon plant exposure to environmental stresses (*e.g*. drought or pathogen attack) or treatments with chemicals disrupting wall biosynthesis (*e.g*. isoxaben). These wall modifications have a direct effect on cell wall integrity (CWI) and can initiate molecular adaptive mechanisms like cell wall composition remodeling and defensive responses activation (12, 18–21). CWI alteration also occurs in plants impaired in or overexpressing cell wall-related genes. Some of these plants/mutants show altered disease resistance phenotypes that were initially associated with the mis-adaptation of pathogens to overcome the modified wall structures of these genotypes (5, 7, 22–27). However, activation of defensive pathways take place in the majority of these mutants/overexpressing lines with wall alterations (5, 7, 22–27). For instance, impairment of cellulose synthesis by inactivating cellulose synthase subunits, as it occurs in *Arabidopsis thaliana irregular xylem mutants* (*irx1, irx3* and *irx5*), or for primary cell walls, as it takes place in *Arabidopsis isoxaben resistant (ixr1)/constitutive expression of VSP (cev1*) mutant, results in constitutive activation of some canonical defensive responses and enhanced resistance to different pathogens. For example, *irx1-6* shows enhanced resistance to the necrotrophic fungus *Plectosphaerella cucumerina* and the vascular bacterium *Ralstonia pseudosolanacearum* (former *R. solanacearum;* 28–31). Similarly, alteration of the biosynthesis and/or structure of wall pectins (*e.g*. degree of methyl-esterification) can also affect pathogen resistance (8, 32–37). Moreover, modification of glucuronoxylans and xyloglucans structure also impacts on disease resistance, as it occurs in the *Arabidopsis de-etiolated3 (det3*) mutant that shows enhanced resistance to *P. cucumerina* (38, 39) or in *agb1* (impaired in Gβ subunit of the heterotrimeric G protein), which has reduced xylose content and shows enhanced susceptibility to several pathogens, including *P. cucumerina*, the biotrophic oomycete *Hyaloperonospora arabidopsidis* and the hemibiotrophic bacterium *Pseudomonas syringae* (40–43). Also, modification of the degree of acetylation of wall polysaccharides and of lignin composition affect disease resistance, growth and adaptation to environmental changes of plants (12, 44–46).

Alteration of CWI can initiate the release of DAMPs that regulate plant immune responses in a similar way to those triggered by Microbe-Associated Molecular Patterns (MAMPs) (6, 24, 47). Despite the diversity of glycan structures of plant cell walls, only a limited number of wall-associated DAMPs have been identified so far, including some oligosaccharides structures derived from β-1,3-glucan (callose), cellulose, xyloglucan, mannan and homogalacturonan polysaccharides of plant cell walls (6, 8–10, 12, 47–50). Modification of CWI leads also to developmental phenotype alterations (*e.g*. reduced plant size, biomass or fertility) indicating that cell wall contributes to plant fitness (5, 19, 51–52). Notwithstanding the evidence of the roles of plant cell walls in immunity and fitness, correlations between variations in cell walls carbohydrate moieties composition and specific phenotypes have not been described until recently (12, 41, 53).

We have investigated the specific contribution of plant cell wall to disease resistance by testing the susceptibility to three different pathogens of a large set of *Arabidopsis* cell wall mutants (*cwm*, n = 38). We found that a significant proportion of these mutants (31 of 38, 81.6%) showed altered disease resistance phenotypes supporting a more relevant function of the walls in plant immunity than currently considered. Here we demonstrate, using analyses of glycome profiling of *cwm* walls, that the content of specific wall glycan moieties in *cwm* plants correlates with some of their disease resistance and fitness phenotypes, providing a link between plant cell wall composition and plant development/immunity phenotypes.

## Results

### *Arabidopsis* cell wall composition specifically contributes to disease resistance responses

To determine the specific function of plant cell wall in immunity, we selected two large sets of *Arabidopsis* cell wall mutants (*cwm*) and tested their resistance to three pathogens with different parasitic styles: the necrotrophic fungus *P. cucumerina* (*Pc*), the vascular bacterium *R. pseudosolanacearum* (*Rp*) and the biotrophic oomycete *H. arabidopsidis* (*Ha*). The first set of *cwm* included eighteen previously described mutants, such as *irregular xylem (irx1-6, irx2-1, irx3-1, irx6-1, irx8-1, irx10-1* and *ixr12-1*) and *powdery mildew resistance (pmr5-1* and *pmr6-1*) mutants, and *det3-1, fra3-1, wat1-1, arr6-3, agb1-2, exp1-1, araf1-1, araf2-1* and *ctl2-1* lines (*SI Appendix*, Fig. S1 and S2). The second *cwm* set comprised twenty T-DNA insertional mutants, which were impaired in orthologs of *Zinnia elegans* genes differentially expressed during xylogenesis, a process involving secondary wall biosynthesis (*SI Appendix*, Fig. S1; 22). We evaluated the resistance phenotypes of these 38 *cwm* lines and their wild-type counterparts (Col-0, Ws-0 or La-*er*) upon infection with *Pc*, *Rp* and *Ha*, either by measuring plants macroscopic disease symptoms (for *Rp* and *Pc*) or conidiospore production (for *Ha*). *irx1-6* plants were included as a resistant control (for *Pc, Rp* and *Ha*), and *agb1-1 (Pc* and *Ha*) and *arr6-3 (Rp*) as hypersusceptible controls (20, 29, 31, 39). We found that 31 of the 38 *cwm* lines tested (81.6%) showed, in comparison to wild-type plants, altered resistance responses (mainly enhanced resistance) to at least one of these pathogens: 22 of 38 mutants to *Pc* and *Ha* (57,9%) and 16 of 38 mutants to *Rp* (42.1%; Fig. 1; *SI Appendix*, Fig. S2). Cluster analyses of these phenotypes identified some specific groups of *cwm* mutants with similar disease resistance phenotypes, but also a high diversity of disease resistance phenotypes illustrated by mutants with unique phenotypes (Fig. 1*A*). Remarkably, 14 of the 38 mutants showed enhanced resistance to more than one pathogen and 3 mutants to all three (*fra3* and *det3*, and the previously characterized *irx1-6*; 29, 31) (Fig. 1; *SI Appendix*, Fig. S2). For *Pc*, the levels of enhanced resistance (*e.g. det3-1, fra3-1, at1g70770-1, ago4-1t sag21-1, irx2-1, at5g51890* or *arr6-3*) or susceptibility (*e.g. xcp2-1, crt1-1, araf2-1* and *akk6-2*) found were lower than those of *irx1-6* and *agb1-1* plants, respectively (Fig. 1B; *SI Appendix*, Fig. S2; 29, 40). *Pc* resistance phenotypes were further validated by infection of a representative set of additional alleles of some mutants (*SI Appendix*, Fig. S3). In the analysis of resistance to *Rs*, we identified 6 mutants that showed enhanced susceptibility and higher disease symptoms (*pdf2.1-2, ago4-t, at3g47510-1, sag21-1, miel1-1* and *arr6-3*) and 10 lower symptoms and enhanced resistance (*det3-1, xcp2-1, irx10-1, fra3-1, wat1-1, ctl2-1, irx1-6, acs8-2, irx6-1* and *irx3-1*) than their corresponding wild-type plants (Fig. 1C; *SI Appendix*, Fig. S2). Except for *fra3-1* and *irx10-1*, the enhanced resistance of these 10 mutants was weaker than that of the previously characterized *irx1-6, irx3-1* or *wat1-1* plants, whereas only *pdf2-1-2* plants were as susceptible as the recently described hypersusceptible *arr6-3* plants (20, 29, 54; Fig. 1*C*; *SI Appendix*, Fig. S2). Notably, we found 14 mutants with enhanced resistance to the biotroph *Ha (e.g. det3, xcp2-1, at1g23170-1, fra3-1, irx10-1, fra3-1, at1g70770-1, at3g47510, acs8-2, at4g15160-1, namt1-1, sag21-1, at5g518-1* and *arr6-3*) and 5 lines that were, like *agb1-1*, more susceptible than wild-type plants to this oomycete (*at1g73440-1, wat1-1, at3g47510-1, irx6-1 and spt42*) (Fig. *1D; SI Appendix*, Fig. S2). These data show a relevant function of the cell wall composition in resistance to different types of pathogens, which was not anticipated.

**Figure 1.**
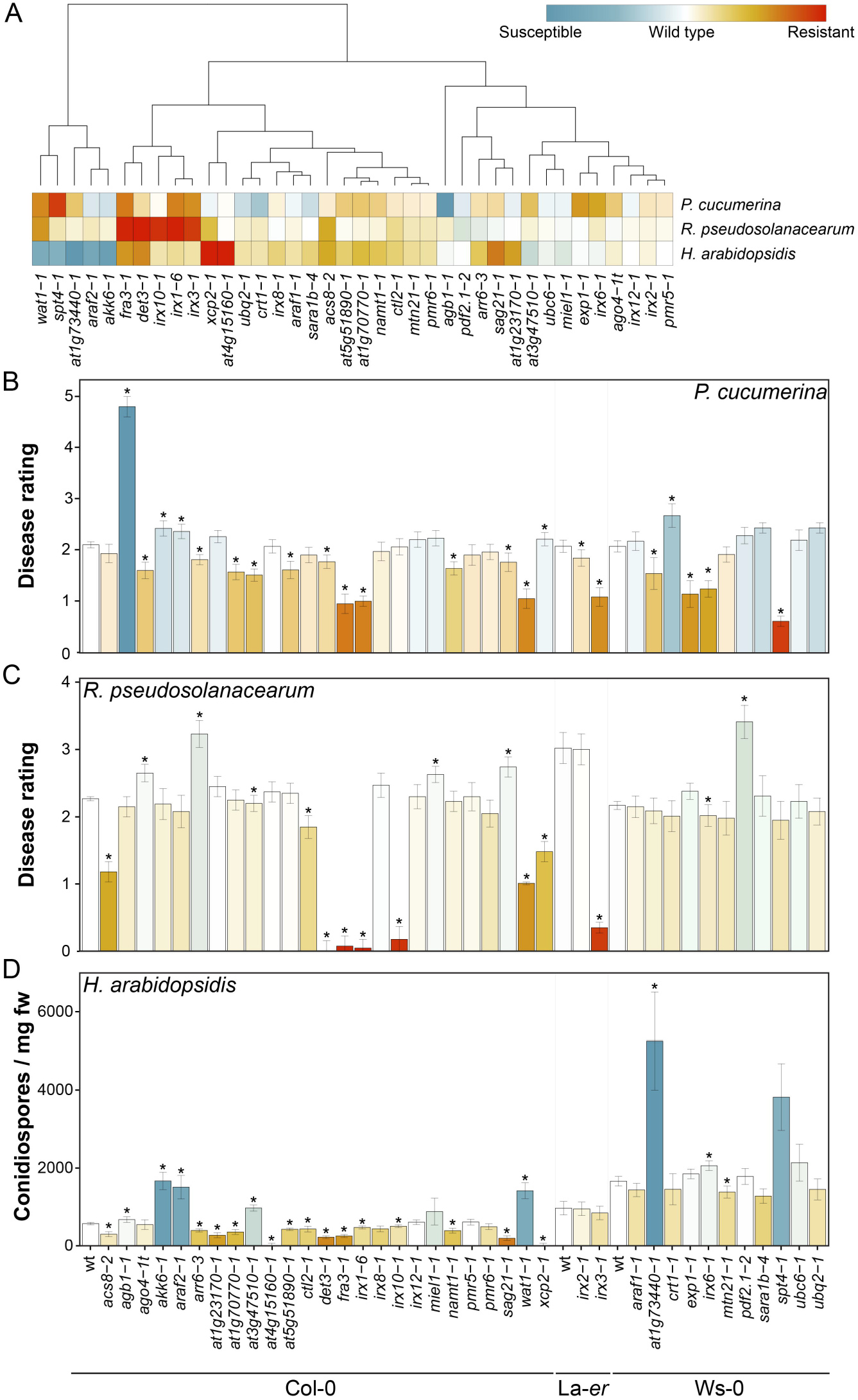
*Arabidopsis* cell wall mutants show alterations of their disease resistance phenotypes in comparison to wild-type plants. (*A*) Clustering of disease resistance phenotypes of *Arabidopsis* cell wall mutants to *P. cucumerina, R. pseudosolanacearum* and *H. arabidopsidis*. Clusters were computed using Euclidean distances using disease resistance indexes relative to wild-type pants (disease rating for *P. cucumerina* and *R. pseudosolanacearum*, number of conidiospores per mg of rosette fresh weight for *H. arabidopsidis*). (*B*) Disease rating (DR; average ± SD) of wild-type (wt) plants (Col-0, La-erand Ws-0. backgrounds) and mutants at 7 days post inoculation (dpi) with the necrotrophic fungus *P. cucumerina*. DR varies from 0 (non-infected plants) to 5 (dead plants). (*C*) DR (average ± SD) of wt and mutants at 8 dpi with bacterium *R. pseudosolanacearum*. DR varies between 0 (no symptoms) and 4 (dead plants). (*D*) Number of conidiospore/mg fresh weight in wt and mutant plants (average ± SD) at 7 dpi with the oomycete *H. arabidopsidis*. Asterisks indicate significant differences compared with wt values (ANOVA non-balanced analysis. Dunnet test p≤0.05). This is one representative experiment of the three performed that gave similar results (n > 10). References and details of *cwm* mutants are listed in SI *Appendix* Fig. S1.

### Enhanced resistance of *cwm* plants to *Pc* and *Rp* negatively impacts plant fitness

The overall developmental phenotypes (*e.g*. rosette size and leaf architecture) of the majority of *cwm* tested did not differ significantly from those of wild-type plants. This is in contrast with the previously described dwarf/altered phenotypes of *irx1-6, irx3-1, fra3-1* or *det3-1* mutants, and that found here for *at5g51890-1 (SI Appendix*, Fig. S4 and see references in Fig. S1). Since disease resistance/developmental growth trade-offs have been described in *Arabidopsis* (4, 55, 56), we selected 18 *cwm* mutants from representative clusters of resistance phenotypes (Fig. 1*A*) and in different ecotype backgrounds, and we measured vegetative (rosette biomass) and reproductive (seed production) traits related to fitness under growth conditions with not limitation of nutrients and water and not infection. Rosette biomass (fresh weight) of 4-week-old plants was, in comparison to wildtype plants, reduced (between 30%-80%) in nine out of 18 *cwm* tested (*det3-1, irx1-6, at4g15160-1, acs8-2, namt1-1, at1g23170-1, fra3-1, irx6-1* and *irx3-1*) and no significant increase in rosette biomass was observed in any of *cwm* lines (Fig. 2*A*). Seed production at the end of reproductive cycle was significantly reduced, in comparison to wild-type plants, in six mutants (*irx10-1, irx1-6, at5g51890-1, fra3-1, irx3-1* and *irx6-1*) and notably increased in two *cwm* lines (*ago4-t1* and *sag21-1*) (Fig. 2*B*). Both fitness traits were negatively affected only in three mutants, *irx1-6, fra3-1* and *irx3-1*, as described previously (29, 52), suggesting that these two traits are decoupled.

**Figure 2.**
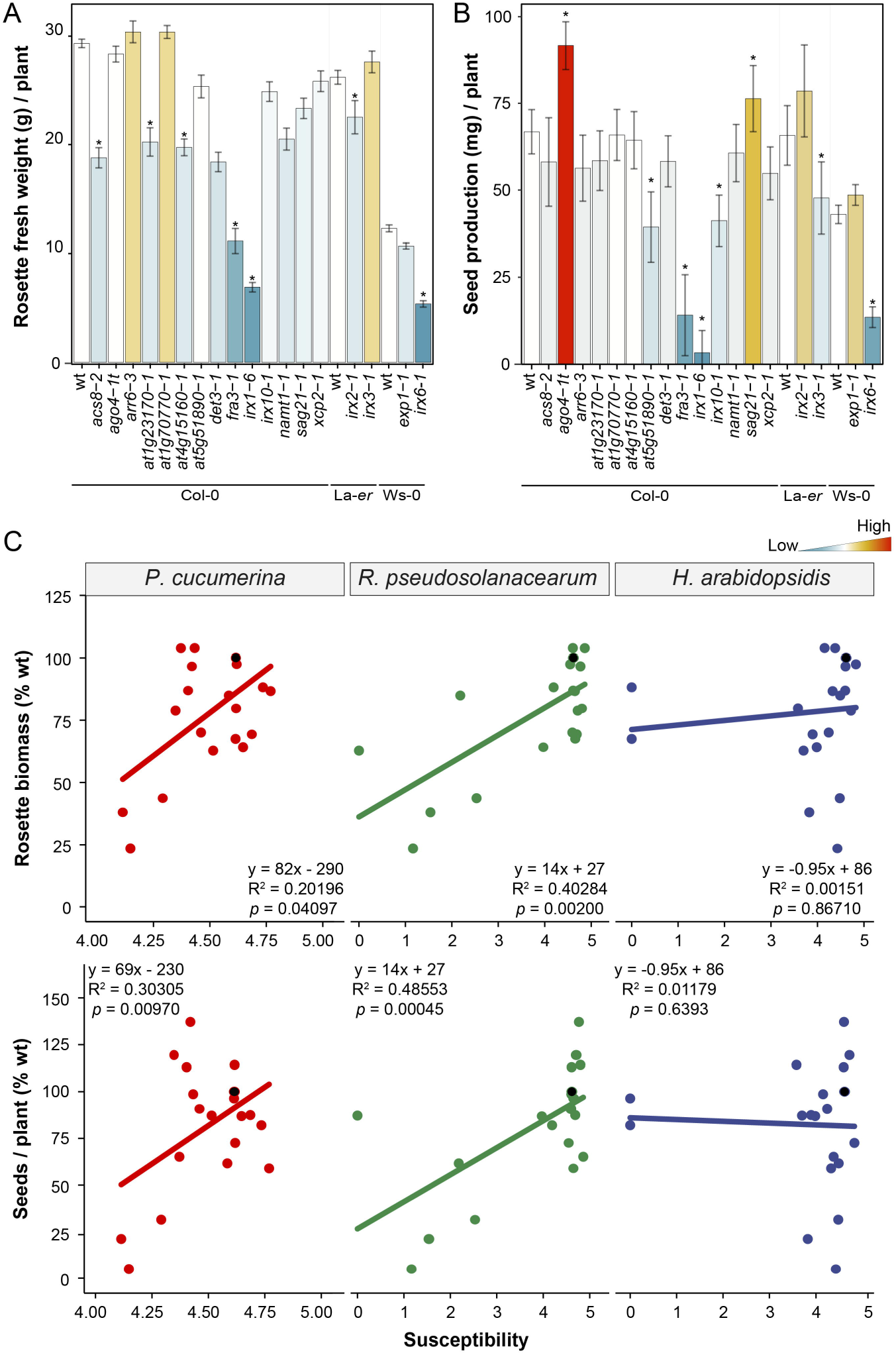
*Arabidopsis* cell wall mutants show associated resistance/fitness trade-offs. (*A*) Rosette fresh weight (average g/plant ± SD) biomass of four-week old mutants and wild-type plants (wt; Col-0, La-er and Ws-0 backgrounds), (*B*) Seed yield (average mg/plant ±SD) at the end of reproductive cycle. Data are the average of 10 plants. Asterisk indicate significant differences compared with wt values (ANOVA non-balanced analysis, Dunnet test p≤0.05). This is one representative experiment of the three performed that gave similar results (n > 10). (*C*) Correlation analysis between biotic stress susceptibility to pathogens (*Pc*, *Rp* and *Ha*) and fitness parameters (seed yield and rosette biomass) of 18 *cwm* mutants and wild-type plants genotypes (3 ecotypes). Average response information of each genotype (dot in the graph) is expressed in relation to that of the reference wild-type plant (black dot, value of 100% at the y-axes). Plants biotic stress susceptibility ratios were log-transformed, and accordingly *x*-axes range from 0 (lower susceptibility) to 5 (greater susceptibility), with the wild-type plants situated at 4.72 = *In* (1 + 100). A linear model was fitted for each combination and correlations determined. Fitted equations, R-squares and p-values are indicated in the insets of graphs. The x-axes of the figures involving *Pc* are enlarged to the 4-5 range for better visualization.

We determined next whether the alteration of fitness of *cwm* was associated with their resistance/susceptibility phenotypes in this subset of 18 *cwm* mutants. Correlation analyses were performed after conversion of DR and fitness data to percentage susceptibility ratios (with respect to each ecotype’s wild-type value) followed by least squares means estimation. A negative correlation was found in *cwm* plants between both rosette biomass and seed production, and resistance to *Pc* (*p* = 0.04097 and R^2^= 020196 for seed yield and *p* = 0.0097 and R^2^ = 0.30305 for biomass) and *Rp* (*p* = 0.002 and R^2^ = 0.40284 for seeds yield and *p* = 0.00045 and R^2^ = 0.48553 for biomass) (Fig. 2*C*). In contrast, a negative association was not found between the resistance phenotype to *Ha* of *cwm* plants and their seeds yield (*p* = 0.8671 and R^2^ = 0.00151) or biomass (*p* = 0.6393 and R^2^ = 0.01179) (Fig. 2*C*). These results indicated that trade-offs between resistance to *Pc/Rp* and plant development exist. Of note, we did not find among the *cwm* mutants with enhanced resistance any with higher seed yield or rosette biomass than wild-type plants (Fig. 2*C*), indicating that *cwm* defensive responses associated to CWI alteration are costly for plant development.

Associations between resistance to *Pc* or *Rp* and tolerance to abiotic stresses (*e.g*. drought, desiccation and salinity) have been reported (29, 54). Accordingly, we quantified the tolerance to desiccation (survival percentage rate upon re-watering after desiccation) of the subset of 18 *cwm*. We found that six of them (*det3-1, irx1-6, fra3-1, irx3-1, irx10-1* and *irx2-1*) were more tolerant to desiccation than the wild-type plants (*SI Appendix*, Fig. S5*A*). Of note, a positive correlation was found between desiccation tolerance of *cwm* plants and disease resistance to either *Pc* (*p* = 0.03141 and R^2^ = 0.22128) or *Rp* (*p* = 7.196 x 10^-6^ and R^2^ = 0.66227), but not to *Ha* (*p* = 0.5254 and R^2^ = 0.02155; *SI Appendix, Fig. S5B*). These results supported previous findings and corroborate that resistance to *Pc/Rp* and desiccation tolerance are linked traits (29, 54).

### Enhanced resistance phenotypes of *cwm* plants are associated with different alterations of their cell wall compositions

We next determined the putative correlations between the observed resistance phenotypes of *cwm* plants and their wall composition (*e.g*. cellulose, neutral sugars and uronic acid content). Of the subset of 18 mutants used in trade-off analyses, only 9 have been previously characterized as cell wall mutants (*SI Appendix, Fig S1*), whereas 9 were putative wall mutants (22). We found, in comparison to wild-type plants, differences in the composition of the walls of the majority of this subset of 18 mutants: *at1g23170-1, at1g70770-1, xcp2-1 ago4-1t* and *irx6-1* had reduced and *acs8-2* increased levels of cellulose, *det3-1* and *irx1-6* possessed less pectic uronic acids, and non-crystalline neutral sugars levels were increased in *arr6-3* and decreased in *at1g70770* and *acs8-2* (*SI Appendix*, Fig. S6). These data confirmed that the majority of 9 putative *cwm* initially selected showed wall alterations. Since these biochemical characterizations of the wall composition of *cwm* plants were not very precise, we narrowed down the collection to a set of 10 mutants, representing six different clusters with different resistance phenotypes, in order to perform a deeper cell wall profiling (Fig. 3*A*). We subjected mutants and wild-type purified cell walls to Fourier-Transform InfraRed (FTIR) spectroscopy that can assign wall polymers and functional groups to different wavenumbers of the FTIR-spectra (57). Differential FTIR-spectra obtained after digital subtraction of the wild-type values from the mutants, showed clearly that *xcp2-1, namt2-1, acs8-2, at1g70770-1, at1g23170-1* and *ago4-1t* were cell wall mutants with biochemical alterations that differ from those observed in the previously characterized *det3-1, irx1-6, irx10-1* and *arr6-3* wall mutants (Fig. 3*B*; 21, 28). Since some of the wavenumbers of the differential FTIR-spectra were associated to lignin components (wavenumbers at 1515, 1630 and 1720 cm^-1^), we determined total lignin content and we found that it was altered in 4 mutants (*det3-1, irx1-6, namt1-1* and *at1g23170-1*) further supporting that these genotypes were mainly affected in secondary wall composition (*SI Appendix*, Fig. S7).

**Figure 3.**
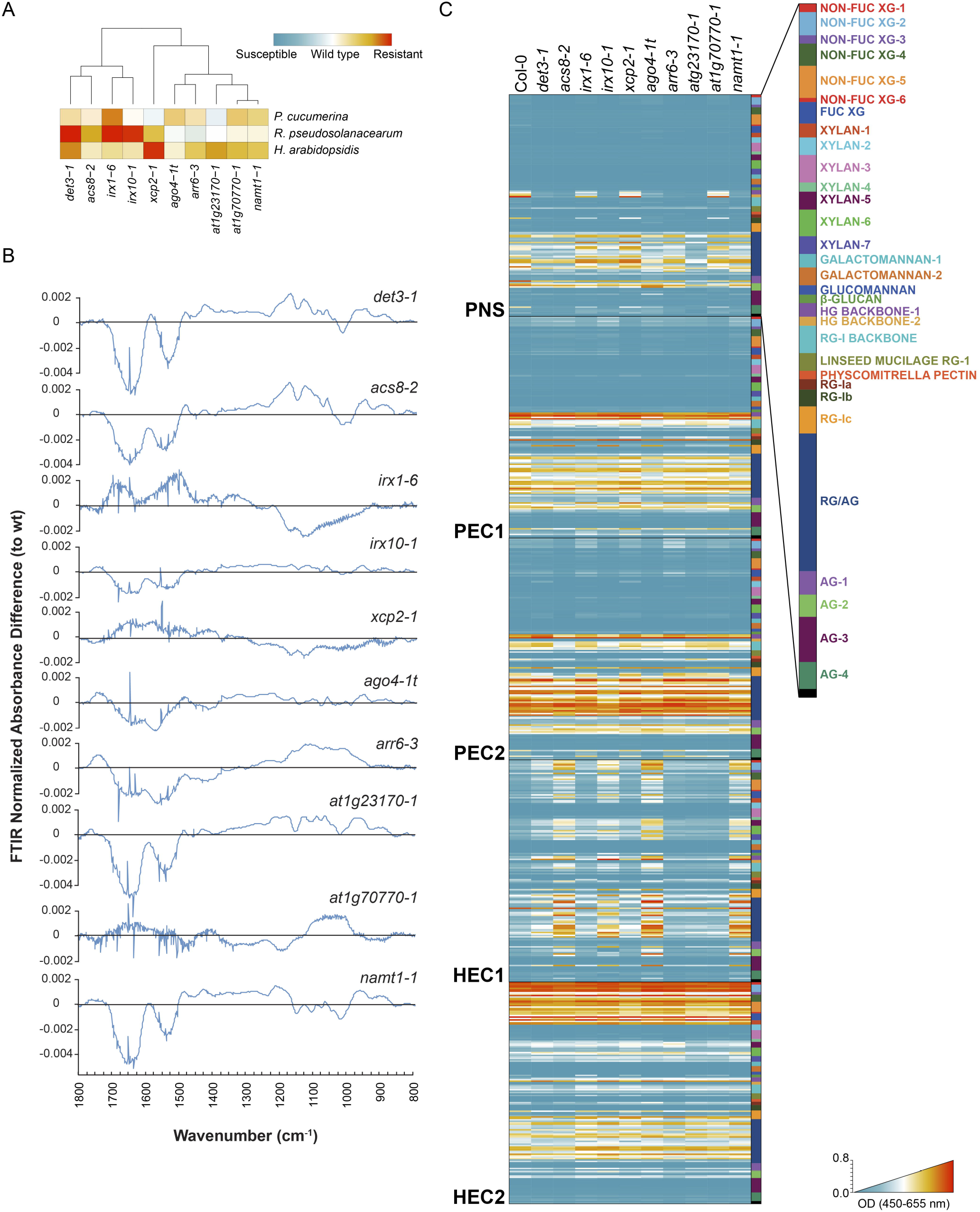
Cell wall analyses by FTIR spectroscopy and glycome profiling of a core set of Arabidopsis cell wall mutants. (*A*) Selection of a core set of representative mutants with different level of disease resistance to *P. cucumerina. R. pseudosolanacearum* and *H. arabidopsidis*. Clusters were computed using Euclidean distances using disease resistance indexes relative to wild-type pants. (*B*) Cell wall FTIR difference spectra of mutants and wild-type plants (Col-0). Black line indicated wild-type values and values over this line differential FTIR spectra in mutants tested. (*C*) Heatmaps of glycome profiling of cell wall extracts (PNS, PEC1, PEC2, HEC1 and HEC2) of *cwm* and wild-type (Col-0) rosette leaves of 25-day-old plants (see SI *Appendix* Table S1 for details). Heatmaps depict antibody binding strength based on optical density (OD) indicated as a color gradient ranging from blue (no binding) to red (strongest binding). The list of monoclonal antibodies used for glycome profiling of each fraction and wall structures recognized by them are indicated in the right panel (see for details SI *Appendix*, Table S1). Data represent average values of two independent experiments (n > 10).

These classical cell wall analyses were complemented with an in-depth compositional characterization of *cwm* wall composition by glycome profiling using a collection of 155 glycan-directed antibodies recognizing diverse cell wall substructures (58, 59). These analyses were carried out on five sequential wall extracts obtained from rosettes of each of plant genotypes: protein and neutral sugars (PNS), and two pectin (PEC1 and PEC2) and two hemicellulose (HEC1 and HEC2) wall extracts, which are known to be enriched in different glycans (58, 59). Glycome profiling confirmed that all the selected mutants showed significant differences in the abundances of some wall glycan epitopes in comparison to wild-type plants (Fig. 3*C*; *SI Appendix*, Table S1), and also corroborated the diversity of wall compositions in the selected wall mutants. The relative abundances of some specific wall epitopes in the PEC1 and PEC2 glycome profiles (*e.g*. fucosylated-xyloglucan) showed opposite patterns in the resistant *cwm* mutants in comparison to those showing wild-type or hypersusceptible phenotypes, suggesting some kind of correlation between specific wall epitopes and disease resistance (*SI Appendix*, Table S1). To test this hypothesis, we performed a model analysis to uncover and generalize the potential relationships between each mutant’s glycomic data (that act as independent or explanatory variables) and disease resistance to *Pc*, *Rp* or *Ha*, as response variables. We used a non-parametric Classification and Regression Tree (CRT) methodology for these analyses, which provides simple and interpretable classification models with almost no statistical assumptions (*SI Appendix*, Fig. S8*A* and Material and Methods). CRT identified a set of antibodies whose reaction values explained, with an estimated cross-validation accuracy between 83.43% and 84.34% (in 10 *cwm* + wild-type genotypes), the resistance phenotypes of *cwm* plants (*SI Appendix*, Fig. S8*B* and Table S2). For example, the abundance of fucosylated-xyloglucans (recognized by CCRC-M106 antibody) correlated with the level of resistance to *Pc* and explained the response phenotypes of 8 out of 11 genotypes tested (Fig. 4*A*). Similarly, CCR5-M5 (detecting an as yet undefined RGI epitope) correlated with the resistance to *Rp* (8/10 *cwm* genotypes), and CCRC-M174 (detecting Galactomannan) and CCRC-M106 (detecting fucosylated-xyloglucans) explained the resistance to *Ha* (8/10 *cwm* genotypes) (*SI Appendix*, Fig. S8*B* and S9). Additional carbohydrate moieties may also contribute to explain mutant’s disease resistance phenotypes but with lower accuracy values (*SI Appendix*, Table S2). To further validate the association of fucosylated-xyloglucan (CCRC-M106) with *Pc* disease resistance phenotype, we performed glycomic analyses with selected antibodies on 3 additional mutants (*pmr5-1, pmr6-1* and *irx8-1*) with resistance to *Pc* similar to wild-type plants (Fig. 1*A*) and on *CA-YDA* plants that overexpress the constitutive active YODA MAP3K and shows enhanced resistance to *Pc* and additional pathogens (60). As predicted by the model, walls of *CA-YDA* plants, but not those of *pmr5-1, pmr6-1* and *irx8-1*, showed an enhanced accumulation of the fucosylated-xyloglucan epitope recognized by CCRC-M106 in comparison to Col-0 cell walls (Fig. 4*B*).

**Figure 4.**
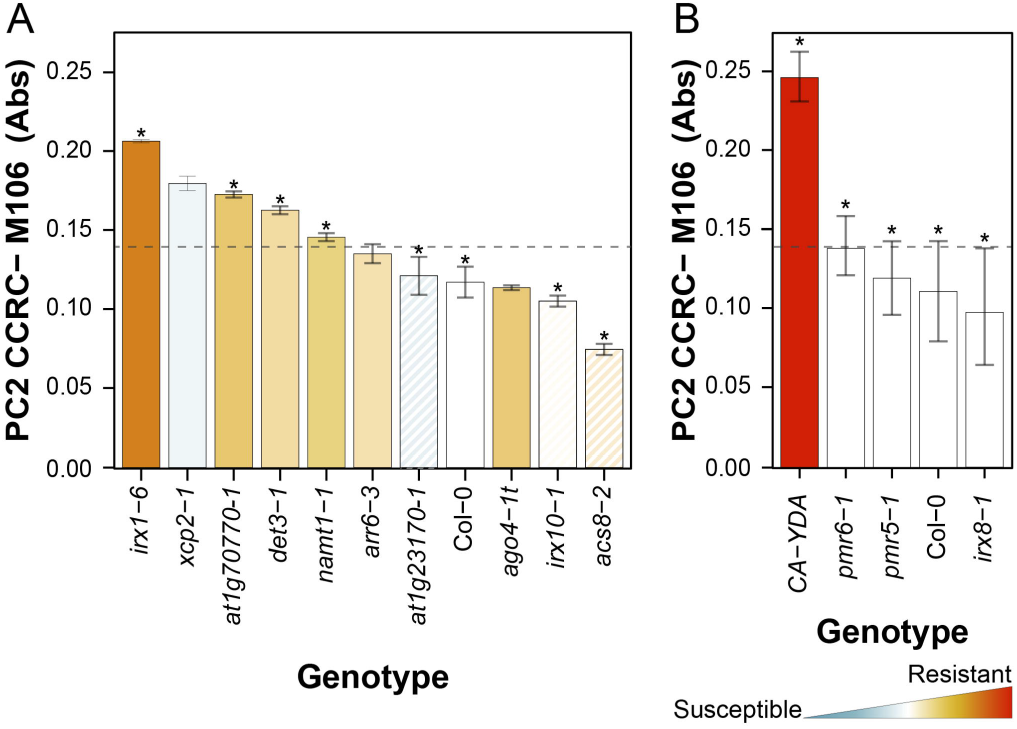
CRT analyses correlate wall composition and disease resistance phenotypes of the *Arabidopsis* cell wall mutants. (*A*). Biological validation of CRT model for resistance to *P. cucumerina* with cell wall mutants from 6 of the cluster analyzed. The absolute value (average ±SD) of the epitope signal detected by CCR-M106 antibody is shown. Columns are colored according to the resistance level of the corresponding mutant, from red (resistant) to blue (susceptible) in comparison with wild-type (wt) level of resistance (white column). Stripped columns mean no significant resistance differences with wt plants (see Fig. 1). The absorbance cut-off value for considering a mutant as resistant or susceptible/wt, as determined by CRT, is indicated by the dotted lines. The *cwm* which follow CRT model are marked with an asterisk, (*B*). Biological validation of CRT model with *pmr5-1, pmr6-1 and irx8-1* mutants, that do not show enhanced resistance to *P. cucumerina* and *CA-YDA* plants, that show enhanced resistance to the fungus. All these mutants follow the absorbance cut-off value predicted by CRT model and accordingly are marked with an asterisk.

Similar CRT analyses were then performed with the fitness parameters (biomass and seed yield, acting as dependent variables) of these 10 *cwm* mutants. Of note, we found a relationship between the reaction signal of some antibodies recognizing some particular carbohydrate moieties and these fitness traits, that explained between 87.31% and 87.62% of the phenotypes: CCRC-M22 (selective for a 6-linked ²-galactan epitope in RGI and arabinogalactan) explained biomass phenotypes, and the levels of epitope detected by CCRC-M175 (galactomannans) and CCRC-M170 (acetylated mannans) correlated with seed yield (*SI Appendix*, Fig. S8*B*, S10 and Table S2). Similarly, we found relationship between tolerance to desiccation and the epitope recognized by the JIM101 antibody (detecting an RGI epitope) that explained 83.16% of the phenotypes (*SI Appendix*, Table S2). Together these data indicate that cell wall compositions of *Arabidopsis* are determinants of plant developmental phenotypes and resistance/tolerance to biotic and abiotic stresses.

### Disease resistance responses of cell wall mutants is not associated to the differential regulation of canonical defensive pathways

The molecular defensive mechanisms underlying the enhanced resistance/susceptibility of the *cwm* lines were further investigated by qRT-PCR determination of the expression of defense genes in non-infected and *Pc*-inoculated plants (1-day post inoculation, dpi). The genes tested are either up-regulated by MAMPs (*e.g. WRKY33, PHI1, CYP81F2* and *PAD3*), CWI alteration (*At1g51890;* 61) or by defensive phytohormone (*PR1, LOX3, PR4, LTP3* and *PDF1.2*, which are gene markers of the salicylic acid (SA), ethylene (ET), jamonic acid (JA), abscisic acid (ABA), and both ET and JA, respectively). We clustered the expression levels of these genes in *cwm* with their resistance phenotypes to identify potential correlations, and only a significant cluster between expression of *LTP3* in non-inoculated *irx1-6* plants and disease resistance to *Pc* was found (*SI Appendix*, Fig. S11 *A* and *B*), as previously described (29). In *Pc* inoculated plants, only one cluster was found associated to *PR1* expression, but it did not explain the *cwm* phenotypes of resistance to any pathogen, since it includes two mutants and wild-type plants (*SI Appendix*, Fig. S11 *A* and *B*). These data clearly indicated that a constitutive expression or enhanced upregulation upon infection of phytohormone-, MAMP-triggered immunity (MTI) or CWI-related genes did not explain the *cwm* enhanced resistance or susceptibility phenotypes.

We have recently shown that pectin wall fractions of Col-0 and *arr6-3* plants contain potential glycan-derived DAMPs that regulate immune responses when applied to Col-0 plants (20). Similar immune responses triggered by elicitor activities have been described in wall extracts of additional *Arabidopsis* wall mutants (12, 44). We found that the PEC1 and PEC2 wall extracts from *cwm* plants, and particularly PEC1 from *at1g23170-1, acs8-2, namt1-1, arr6-3* and *irx10-1*, triggered early immune responses, like Ca^+2^ bursts, upon their application to Arabidopsis *pMAQ* lines expressing the apoaequorin Ca^+2^ sensor protein (20, 47; *SI Appendix*, Fig. S12). These data suggest that these wall extracts might contain additional DAMPs or increased amounts of DAMPs, in comparison to wild-type ones, that might regulate *Arabidopsis* immune responses contributing and explaining the disease resistance of the mutants.

## Discussion

Plant cell walls are important components of both preexisting and inducible plant defense mechanisms against pathogens infection (5, 6, 18, 19). Accordingly, modifications of cell wall composition and structure in some mutants or transgenic lines have been demonstrated to result in the alteration of their resistance phenotypes to different pathogens, including hemibiotrophic (*e.g. P. syringae;* 8, 29, 63) and vascular (*e.g. R. pseudosolanacearum*; 29, 40, 54) bacteria, and necrotrophic (*e.g. P. cucumerina* and *Botrytis cinerea*; 21, 29–32, 36 39, 42, 54, 64) or biotrophic fungi (*e.g. Erysiphe* sp. 28, 65–68). For example, 15 of the wall mutants analyzed in this study have been previously described to show differential disease resistance phenotypes to one or two pathogens, and in a few cases, like *irx1-6, agb1-1* and *arr6-3* used as controls, their resistance to three or more pathogens have been determined (Fig. 1, *SI Appendix Fig. S1*). Despite these previous data, specific correlations between wall composition/structure and the resistance phenotypes and/or immune responses of these plant genotypes have not been described.

Here, we have determined the contribution of *Arabidopsis* cell walls to disease resistance responses against three pathogens with very different life-styles by selecting a large set of cell wall mutants, that includes well characterized and putative wall mutants (22). Given the different molecular bases of *Arabidopsis* resistance to the three pathogens tested (*Pc*, *Rp* and *Ha*; 20, 29, 39, 40, 54, 62) and the putative diversity of cell wall alterations in the mutants tested, we initially anticipated that we could obtain a global view of *Arabidopsis* cell walls contribution to resistance. Notably, we found that 81.6% of the cell wall mutants tested (31 of 38) showed, in comparison to wild-type plants, differential phenotypes (enhanced resistance (mainly) or susceptibility (in a few cases)) to at least one of the three pathogens tested. Of note, we have identified different clusters containing one or several mutants with specific phenotypes (*e.g*. from enhanced resistance to the three pathogens to specific resistance to one pathogen; Fig. 1*A*). These data support the diverse and significant functions of cell walls on plant disease resistance responses to vascular and necrotrophic pathogens, as previously described (20, 29, 54). Our data also identified an unanticipated contribution of cell walls to plant resistance to biotrophic oomycetes, like *H. arabidopsidis* (Fig. 1*C* and *SI Appendix*, Fig. S2), that is in line with the described wall function in *Arabidopsis* resistance to biotrophic fungi causing powdery mildew diseases (65–69). The proportion of mutants with differential disease resistance phenotypes identified in this screening is several orders of magnitude higher than the expected proportion that would be obtained in blind, unbiased disease resistance screenings using T-DNA or chemically mutagenized plant populations. It can be anticipated that a similar proportion of genotypes to that obtained here would be found if a biased screening was performed with *Arabidopsis* mutants impaired in known components of key defensive pathways, like phytohormone signaling or MAMP-triggered immunity (70, 71). Therefore, the data obtained here with the set of *cwm* plants strongly support the relevant function of cell walls in plant immunity and disease resistance to different pathogens.

The genetic and molecular basis of *Arabidopsis* resistance to the three pathogens analyzed here differ significantly: (I) plant resistance to *H. arabidopsidis* mainly depends on activation of effector-triggered immunity (ETI) and of the SA pathway; (II) disease resistance to *R. pseudosolanacearum* is mediated by ET and just a few examples of ETI responses have been described; and (III) Arabidopsis resistance to necrotrophic fungi, including *P. cucumerina*, has been shown to depend on several hormone signaling (mainly ET and JA, but also SA) and on the synthesis of Tryptophan-derived metabolites (like camalexin and indole glucosinolates), and few examples of ETI-mediated resistance have been described so far (72, 73). This lack of source of resistance genes for controlling *R. pseudosolanacearum* and necrotrophic fungi, like *P. cucumerina* or *B. cinerea*, might explain the strong incidence of the diseases caused by these two types of pathogens in crops and the associated yield losses, since breeding programs have not been effective in selecting traits conferring enhanced resistant to these pathogens (74, 75). This contrasts with the effectiveness of breeding programs in controlling biotrophic pathogens, like the oomycete (*e.g. H. arabidopsidis*) causing downy mildews or fungi (*e.g. Erysiphe* sp.) causing powdery mildew diseases (73, 76). Our data indicate that modification of cell wall integrity could be, initially, an effective strategy in the control of diseases caused by necrotrophic and vascular pathogens and therefore that a genomic-assisted breeding selection of CWI-associated traits could be used in breeding programs, as suggested previously for other crop traits like biomass digestibility (77). However, modification of cell wall composition and structure usually results in alterations of plant developmental phenotypes (*e.g*. reduced plant size, biomass or fertility) that impact on fitness (19, 78). In line with these previous data, we describe here a negative correlation between fitness parameters, like rosette biomass and seed production, of the cell wall mutants tested, and their enhanced resistance to vascular and necrotrophic pathogens (Fig. 2*C*). These trade-offs associated to increased resistance to these pathogens are also probably hampering the selection of crops traits conferring improved resistance. In contrast, we have not found associated trade-offs to the enhanced resistance to the biotroph *H. arabidopsidis* (Fig. 2*C*), indicating that some wall-associated traits identified here might be of interest for improving resistance to biotrophic pathogens.

Plant cell walls (primary and secondary) are complex and dynamic structures composed mainly of carbohydrate-based polymers of differing monosaccharide and glycosyl-linkage compositions (13). Among the mutants included in our analysis we selected 17 previously described as showing a great diversity of wall alterations, like reduction/alteration of the content/decorations of cellulose (*irx1-6, irx3-1, irx6-1, irx2-1* or *ctl2*), xylan (*irx10-1*), glucuronoxylan (*irx8-1*), pectin (*pmr6-1, pmr5-1, arr6-3* and *irx8-1*), xyloglucans (*agb1-1*) or lignin (*irx12-1*; see SI Appendix Fig. S1 for references), or impairment in glycan transport or *in muro* biosynthesis of wall components (*e.g. ctrl1-1, det3-1, wat1-1* and *fra3-1*; see *SI Appendix* Fig. S1 for references). We also tested 21 putative cell wall mutants (SI Appendix Fig. S1-S2), including some that have been recently characterized as wall mutants (*e.g. araf2-1* and *araf1-1* impaired in arabinan-containing pectins or *xcp2-1* and *arr6-3*), and seven mutants whose wall alteration have been demonstrated for the first time here (Fig 3. and *SI Appendix*, Fig. S6, S7 and Table S1). These data corroborate that the majority of genotypes initially selected are *bona fide Arabidopsis* wall mutants. Cell wall modifications identified by FTIR-spectroscopy or biochemical analyses in the mutants from the 6 phenotypic clusters selected (Fig *3A* and *SI Appendix*, Fig. S6 and S7) were not precise enough to find specific associations between wall composition and disease resistance phenotypes. Chemically extracted cell walls fractions (*e.g*. PEC1 and PEC2) contain mixtures of carbohydrate moieties derived from various polymer classes and can be enriched in certain carbohydrates detectable by glycome profiling. We show here that glycome profiling analysis of these extracts provide a more precise picture of wall modifications impacting disease resistance. Our data also show that mathematical modelling by CRT of glycome profiling of plant genotypes provides detailed and biologically consistent links between cell wall composition and disease resistance/fitness phenotypes, as it has been previously reported for the determination of cell wall digestibility of plant genotypes biomass (12, 79, 80).

The CRT algorithm used here allows both the identification of variables (cell-wall components recognized by some antibodies) and the definition of cut-points on these variables, separating mathematically in different branches and nodes the genotypes belonging to different phenotypic classes (best, equal or worse than wild-type phenotypes). Since CRT is based on binary branching, it obtains more pure or homogenous nodes (in terms of their class composition) in contrast to other supervised classification methods (*e.g*. linear discriminant analysis, logistic regression, random forest or classification trees). Using CRT, we have identified significant epitope associations explaining as much as 84.34% of the disease resistance phenotypes tested (Fig. 4*B*, *SI Appendix* Fig. S8*B* and Table S2). Remarkably, the abundance of fucosylated-xyloglucan (CCRC-M106), RGIa (CCRC-M5) and galactomannan (CCRC-M174)/fucosylated-xyloglucan (CCRC-M106) in the cell walls of the mutants correlated with the level of resistance to *P. cucumerina, R. pseudosolanacearum* and *H. arabidopsidis*, respectively (Fig. 4*A,B*, SI Appendix Fig S9 and S10). The relevance of xyloglucan and xylose content in Arabidopsis disease resistance to *P. cucumerina* has been previously described (39), and is further supported here by the validation of the content of wall epitope recognized by the CCRC-M106 antibody, which is enhanced in the *P. cucumerina* resistant *CA-YDA* plants (Fig. 4*B*; 60). In contrast, the contribution of galactomannan and RGIa/fucosylated-xyloglucans to *R. pseudosolanacearum* and *H. arabidopsidis* resistance, respectively, has not been previously described. Notably, we have also demonstrated here that the level of a 6-linked β-galactan epitope present in RGI and arabinogalactans (CCRC-M22) and of acetylated-mannan/galactomannan (CCRC-M170/CCRC-M175) correlated with rosette biomass and seed production, respectively, indicating that wall composition can also determine plant fitness. These data are in accordance with previous results showing that high-density quantitative glycan microarrays, used in conjunction with association mapping, can detect pertinent variations related to plant cell wall genetics (12, 79, 80). Since the specificity of the carbohydrate moieties recognized by some of the antibodies identified here has not been fully established yet, in contrast to other antibodies of the glycomics collection (59), it cannot be excluded that disease resistance/fitness traits could be associated to other types of wall epitopes than those described here.

Various hypotheses have been put forward to explain why modification of cell wall composition often appears to enhance rather than reduce plant disease resistance (7, 12). These hypotheses include the modification of wall composition that is not hydrolyzed by cell wall degrading enzymes secreted by pathogens, but also the activation of immune responses upon the release of elicitor-active molecules (DAMPs) from incorrectly assembled plant cell walls that will be recognized by plant PRRs triggering uncharacterized immune responses. Our data support that some particular immune pathways are differentially regulated in some of the mutants, but their expression patterns do not explain their disease resistance phenotypes (*SI Appendix*, Fig. S11). Notably, we also show that cell wall fractions of some *cwm* plants might contain additional DAMPs or enhanced levels of DAMPs in comparison to wild-type fractions that might trigger immune responses (*SI Appendix*, Fig. S12), as it has been described recently to occur in other *Arabidopsis* wall mutants (12, 20, 44). This DAMP-triggered immunity together with the canonical-immune pathways, that might be constitutively expressed or primed for stronger activation upon pathogen infection in some cell wall mutants, would contribute to regulate their immune responses and disease resistance phenotypes. The characterization of these *cwm* defensive responses, and the wall DAMPs and PRRs involved in their activation, deserves further attention to characterize these novel wall-associated immune responses.

## Experimental Procedures

### Plant materials and growth conditions

*Arabidopsis* genotypes used in this study and oligonucleotides used for T-DNA insertional mutant characterization are listed in *SI Appendix* (Fig. S1 and Table S3). For plants used in *R. pseudosolanacearum* assays, seeds were germinated on MS medium (82), and then grown in Jiffy pots (Jiffy, http://www.jiffygroup.com) in a chamber at 22°C, with a 9 h light period and a light intensity of 200 μmol m^-2^ sec^-1^ and 50% relative humidity. Plants used in *P. cucumerina* and *H. arabidopsidis* disease resistance and fitness experiments were grown on soil in a growth chamber as described (20). Plant rosette biomass was determined on four-weeks old plants (n = 10) and seeds were harvested at 8 weeks after plants (n = 10) completed their vegetative cycle. Experiments were repeated at least three times with similar results.

### Pathogens growth conditions and plant infections

*P. cucumerina* (BMM strain) and *H. arabidopsidis* (isolates Noco, Emwa1 and Cala) were grown as described (20). *R. pseudosolanacearum* (strains GMI1000 and RD15) were grown at 28°C on BG medium (83). For *P. cucumerina* infection, three-weeks old plants (n > 10) were sprayed with a suspension spore (4×10^6^ spores/ml) of virulent *Pc*BMM isolate and progression of the infection was followed by visual evaluation of disease rating at different days post inoculation (dpi) as average disease rating (DR, from 0 to 5) was scored: 0 = no symptoms; 1 = plant with some necrotic spots; 2 = one or two necrotic leaves; 3 = three or more leaves showing necrosis; 4 = more than half of the plant showing profuse necrosis; and 5 = decayed/dead plant (39). For *H. arabidopsidis* assays, 12-day-old plants (n > 20) were sprayed with a conidiospore suspension (2×10^4^ spores/ml) of virulent isolates (Noco2, Emwa1 and Cala for plants in Col-0, La-*er* and Ws-0 backgrounds, respectively), and then plants were incubated under short day conditions (10 h illumination) for 7 days, and the aerial parts of all plants were harvested, released conidiospores, after shaking in water, were counted and their average/mg rosettes fresh weight determined (47). For *R. pseudosolanacearum* infections, roots of four-weeks old plants (n > 10) were dipped into a bacterial suspension (5×10^7^ cfu/ml) of virulent strains GMI1000 (for Col-0 and La-*er*) or RD15 (for Ws-0). Following inoculation, plants were transferred to a growth chamber under the following conditions: 12 h photoperiod, 27°C, 80% relative humidity. Average DR was scored in leaves as follows: 0 = no symptoms, 1 = 25% wilted leaves, 2 = 50% wilted leaves, 3 = 75% wilted leaves, 4 = 100% wilted leaves (dead plant; 62). All pathogens resistance assays were repeated at least three times.

### Plant cell wall purification, fractionation and analyses

Cell walls (Alcohol Insoluble Residues; AIR) were prepared from 25-day-old Arabidopsis plants according to (81), and non-cellulosic fraction, uronic acid and crystalline cellulose and lignin contents were determined as previously described (60, 84). Fourier-transform infrared (FTIR) spectroscopy determination was done with discs prepared from mixtures of purified AIR and KBr (1:100, w:w) using a Graseby-Specac press. FTIR-spectra were recorded and analyzed as described (85). Lignin-like material was quantified by the Klason gravimetric method with minor modifications (86). AIR fractions were subjected to sequential chemical extraction with increasingly harsh reagents in order to isolate fractions enriched in various cell wall components as previously described: Protein and Neutral Sugar (PNS) fraction, pectic fractions (PEC1 and PEC2), and hemicellulosic fractions (HEC1 and HEC2; 20, 81). Glycome profiling of the cell wall fractions was carried by ELISA using a toolkit of plant cell wall-directed monoclonal antibodies as previously described (see *SI Appendix* and 58, 59). Monoclonal antibodies are annotated in a database at www.WallMabdb.net; specific links in Table S1).

### Gene expression analyses

Plants were mock-treated or *Pc*BMM-inoculated, and rosettes (n > 25) were collected at 1 dpi (four biological replicates). Total RNA was extracted using the RNeasy Mini Kit (Qiagen) as reported previously (39) and quantitative real-time PCR amplification (qRT-PCR) and detection were carried out as previously described (47). Oligonucleotides used for gene expression are detailed on Table S4. The expression levels of each gene, relative to *UBC21 (AT5G25760*) expression, were determined using the Pfaffl method (87).

### Clustering and statistical analyses (see also *SI Appendix*)

Heatmaps and cluster aggrupation (Figure *1A*, Figure *3A,C* and Figure S11) were calculated using “gplots” R package version 3.0.3 (88). Clusters in Figure *1A* and Figure *3A* were computed using Euclidean distances using disease resistance indexes relative to wild-type plants (disease rating for *P. cucumerina* and *R. pseudosolanacearum*, number of conidiospores per mg of rosette fresh weight for *H. arabidopsidis*). Clusters in Figure SI11 were computed using Euclidean distances for absolute gene expression levels and disease indexes.

ANOVA models were fitted for each of the response variable (resistance to *Pc*, *Rp*, *Ha*, and biomass and seeds yield, and desiccation tolerance; Fig. 1, Fig. *2A,B*, *SI Appendix*, Fig. S5*A*) and each ecotype (Col-0, Ws-0, or La-*er*). Least squares means (LS means) of these models were then obtained, providing a single estimation of the average response level (*e.g*. mean disease rating for both *Pc* and *Rp*, spore/mg for *Ha*, seed yield in mg and rosette fresh weight in mg, and survival rate after desiccation) for each genotype. Afterwards, correlation analyses (*SI Appendix*, Fig. S13) between biotic resistance and fitness features/desiccation were obtained by determining the ratio of each genotype LS mean to that of the corresponding wild-type ecotype for each response variable (*e.g*. percentage susceptibility levels with respect to wild-type plants). A logarithmic model was fitted for each combination of the biotic susceptibility ratios with the fitness and abiotic susceptibility ratios to analyze their correlations (see Fig. 2*C* and *SI Appendix* Fig. S5*B* for the fitted equations, R-squares and *p*-values). For more details, see *SI Appendix* Material and Methods.

CRT predictive classification model (*SI Appendix*, Fig. S8) correlating wall composition with disease resistance and fitness phenotypes was done by first performing a paired comparison analysis to assign Arabidopsis wild-type and *cwm* mutants genotypes into a class (*e.g*. a categorical valuation), which represent its status compared to wild-type: plants with a similar performance (class equal or E), significantly better (B) or significantly worse (W) than wild-type ones. The CRT method was then applied to link this class status to glycomics data. To avoid overfitting the data, the tree-growing process of each CRT model was limited to a single binary branching, to select a single antibody and its optimal cut-off point. The actual predictive capability or accuracy of the resulting classification tree models is evaluated as the percentage of correctly classified genotypes obtained through a 10-fold cross-validation process replicated 100 times. The correlation and paired comparison analyses were implemented using the SAS software (*glm* and *corr* procedures), while the CRT classification model fit and validation were implemented using Python (*scikit-learn* library). See *SI Appendix* Material and Methods for further details.

## ACKNOWLEDGEMENTS

This work was supported by the Spanish Ministry of Economy and Competitiveness (MINECO) grants BIO2015-64077-R and RTI2018-096975-B-I00 of Spanish Ministry of Science, Innovation and Universities (MICIU) to A.M., by the French National Agency for Research ANR□07□GPLA□014 grant to D.G. This work has been also financially supported by the Severo Ochoa Program for Centers of Excellence in R&D from the Agencia Estatal de Investigación of Spain (grant SEV-2016-0672 (2017-2021) to the CBGP). In the frame of this program H.M. was a postdoctoral fellow and he was also supported by IEF grant (SignWALLINg-624721) from the European Union. E.M. was a Juan de la Cierva Postdoctoral Fellow from MINECO and L.B. FPI fellow of MICIU. The generation of the CCRC-series of plant cell glycan-directed monoclonal antibodies used in this work was supported by the US National Science Foundation (DBI-0421683 and IOS 0923992) to M.G.H. We thank Yves Marco and Philippe Ranocha for his help with *cwm* mutant selection and screening with *R. pseudosolanacearum*. We thank the Molina laboratory members for useful discussion and comments on the manuscript, Javier Paz-Ares (Centro Nacional de Biotecnología, Spain) for their fresh ideas and interpretations about our challenging data, and Fernando García-Arenal (CBGP, Spain) for critical reading of the manuscript.

## Appendix SI Information

- **Figure S1. *Arabidopsis thaliana* mutants used in the disease resistance analyses.**
- **Figure S2. Disease resistance analysis of Arabidopsis cell wall mutant**.
- **Figure S3. Disease rating (DR) of *cwm* second alleles inoculated with *P. cucumerina* BMM.**
- **Figure S4. Developmental phenotypes of three-week old plants of *cwm* and wild-type ecotypes (Col-0, Ws-0 and La-*er*).**
- **Figure S5. Desiccation tolerance of wild-type plants and cell wall mutants.**
- **Figure S6. Cell wall biochemical composition of *Arabidopsis cwm* plants.**
- **Figure S7. Total lignin content of *Arabidopsis cwm* plants**.
- **Figure S8. Predictive CRT model correlating wall composition and disease**
- **resistance/fitness/desiccation phenotypes of *Arabidopsis* cell wall mutants**.
- **Figure S9. Predictive CRT model correlating wall composition and disease resistance/fitness phenotypes of *Arabidopsis* cell wall mutants to *H. arabidopsidis andR. pseudosolanacearum***.
- **Figure S10. Predictive CRT model correlating wall composition and fitness phenotypes of *Arabidopsis* cell wall mutants**.
- **Figure S11. Clustering of the expression pattern of canonical defensive genes in wild-type plants and *cwm* mutants and their disease resistance phenotypes**.
- **Figure S12. Cell wall derived extracts from pectin fractions (PEC1 and PEC2) of *cwm* plants trigger Ca^2+^ elevations.**
- **Figure S13. Schema of the mathematical analyses performed to generate the data of Figures 2C, Fig. 4, Fig. S5B, Fig S8-S10.**
- **Table S1: Glycome profiling of the five different cell wall fractions obtained from different genotypes from the core set of cell wall mutants and Columbia-0 (Col-0) wild-type plants.**
- **Table S2: Summary of the results from the performed Classification and Regression Tree (CRT) analysis.**
- **Table S3: Oligonucleotides used for T-DNA insertional mutant characterization.**
- **Table S4: Oligonucleotides used for gene expression analysis.**

