## Supplementary material for "*Arabidopsis* cell wall composition determines disease resistance specificity and fitness": SI Appendix

**This PDF file includes:**

Supplementary text
Figures S1 to S13
Tables S2 to S4
SI References 1-44

**Other supplementary materials for this manuscript include the following:**

Supplementary Table S1

SI *Appendix*

**Supplementary Material and Methods**

***Determination of $Ca^{+2}$ burst upon treatment with cell wall fractions***

Eight-day-old liquid-grown apoaequorin-expressing seedlings (Col-$0^{AEQ}$) were used for $Ca^{2+}{}_{cyt}$ upon cell wall fractions treatment. Col-$0^{AEQ}$ plants (were grown in 24-well plates (~10 seedlings per well) under long day conditions (16 h of light) at 20-22ºC in liquid MS medium as described (21). Then, they were placed individually in 96-well plates in coelenterazine (PJK GmbH) and water and were incubated overnight in the dark. Luminescence was recorded with a Varioskan Flash Multimode Reader (Thermo Scientific) as described, after treatment with cell wall extracts (PCI and PCII) from wild-type plants and cell wall mutants (1).

***Plant tolerance to desiccation assays***

Three-week-old soil-grown plants (n=8) were restricted completely from irrigation for 21 days and then wilted plants were re-watered for 7 days and the number of plants that survived to the stress and recovered their developmental phenotype was scored. These experiments were performed four times.

***Mathematical Modelling***

The description of the mathematical analyses performed is shown in the schema of *SI Appendix*, Fig. S13, and can be divided into two interrelated tasks that share a set of analytical steps. These common analyses were initiated with the experimental data on susceptibility to pathogens, fitness and tolerance to desiccation of *cwm* and wild-type plants (Fig. 1 and Fig. 2A, B, and *SI Appendix* Fig. S2 and Fig. S5A). For each of these 6 response variables (resistance to 3 pathogens, 2 fitness parameters and plant desiccation tolerance) a different two-way ANOVA model was fit for each ecotype (Col-0, Ws-0, or La-*er*), considering the genotype as the factor/effect of interest, but also introducing a second factor/effect to allow collecting together the data from a number of similar experiments (2 independent experiments in the case of pathogen resistance, 3 for fitness and 3 for tolerance to desiccation). No interaction was initially considered between both factors/effects, rendering the experiment effect as a block. These initial analyses aim to confirm the known significance of the genotype factor/effect, and to estimate the least squares means (or LS means) of each response in each genotype. These LS means provide a single estimation of the average response level (i.e. mean disease rating for both *Pc* and *Rp*, mean spores/mg for *Ha*, mean seed yield in mg, mean rosette fresh weight/biomass in g, and mean survival rate (%) after desiccation) for each genotype under consideration, controlling the effect of the experimental differences due to the unbalanced nature of the designs.

The correlation analysis between mean biotic stress resistance, on one side, and mean fitness and desiccation tolerance, on the other side, was then performed on this intermediate set of 21 genotypes. To this aim, the percentage ratio of each genotype LS mean to that of the corresponding ecotype WT was obtained for each response variable. These susceptibility ratios allowed expressing the average response information of each genotype in relation to that of the reference WT, as well as bringing the information of different ecotypes to a similar scale. A linear model was then fit for each combination of the log-transformed biotic susceptibility ratios with the fitness and abiotic susceptibility ratios to analyze their correlations (see Fig. 2C,D and SI *Appendix*, Fig. SI5; for the fitted equations, *R*-squares and *p*-values; SI Appendix Figure S13). As a consequence of the logarithmic transformation of the biotic susceptibility ratios, the *x*-axes in these figures range between 0 (smaller susceptibility) to 5 (greater susceptibility). The SAS software has been used to implement the previous analyses (*glm* and *corr* procedures).

Next, a paired comparisons analysis on the mentioned LS means was done, and experimental data on glycomic response of cell wall mutants' fractions (Fig. 3C and SI *Appendix*, Table S1) was then incorporated to obtain predictive classification models (correlating wall composition with disease resistance and fitness phenotypes. More specifically, on the basis of the *p*-values of the one-sided Dunnett tests ($\alpha = 0.05$), each genotype was assigned a class (i.e. a categorical valuation) for each response, which represent its status in such feature compared to the WT: either a plant has a similar performance to the WT (class *E*), or the plant performs significantly better than the WT (class *B*) or the plant performs significantly worse than the WT (class *W*). A supervised classification methodology was then applied in order to correlate wall composition with biotic stress resistance, fitness, and desiccation tolerance phenotypes as represented by the mentioned classes *E*, *B* and *W*. The available data to describe wall composition was given by 32 glycomic measurements of a set of 155 antibodies (for each cell-wall fraction) performed for 11 different genotypes of the Col-0 ecotype (Fig. 3) Thus, the objective of the present analysis was to uncover and generalize the potential relationships between the set of glycomic responses (that therefore act as independent or explanatory variables) and the classes that describe the performance of the genotypes (acting as dependent variables), for each of the studied features, to explain the different phenotypical status of the genotypes in the core set, and to generalize such patterns in order to provide a mathematical model exhibiting an adequate predictive accuracy.

Many statistical and machine learning techniques are available to fit supervised classification models, as for instance linear discriminant analysis, logistic regression, random forest or classification trees. In the present analysis we have chosen this last technique (classification trees) since it constitutes a well-known, standard classification methodology with almost no statistical assumptions that provides interpretability of the resulting models, automatic independent variables selection, and an adequate predictive capability for the purposes of the present study. More specifically, the particular classification tree algorithm applied, known as CRT (or CART, from Classification and Regression Tree), is a non-parametric technique that makes no assumptions on the distribution of the data and the variability and balance between classes. The bases of the CRT algorithm are both the identification of independent variables (in this case, cell-wall antibodies) and the definition of cut-points on these variables' values that allow separating in different branches of a tree the instances belonging to different classes. In this way, departing from an initial *root* node containing all the instances of the training sample, CRT tries to obtain purer or more homogenous nodes (in terms of their class composition) by computing the reduction of the node's impurity allowed by each of the available independent variables. Given a set of $n$ classes, each appearing in a given node with a relative frequency $f_i$, $i = 1,\ldots,n$, CRT computes the node's impurity through the Gini Index

$$I_G = \sum_{i=1}^{n} f_i(1 - f_i).$$

Thus, from a given node, CRT selects both the variable and its cut-point that allow the greatest impurity reduction from the "father" node to the "child" nodes. CRT is based on binary branching, i.e. only two child nodes are produced from each father node being split. The main inconvenient of the CRT methodology is its tendency to produce overfit, i.e. to provide classification models that generalize not only the right, essential patterns or relationships between the explanatory variables and the classes, but also the potential noise that the specific data used to construct (or train) the model may contain. When overfit is present, the obtained model tends to provide excellent results on the training sample, but quite worse results on unseen instances not used in the construction of the trees, what is usually called a validation or test sample. To avoid overfit, we force the tree growth process to stop after the first split of the root node, and at the same time we carry out a demanding cross-validation process in order to get a precise estimation of the actual model's accuracy. Therefore, the obtained tree is composed of just the root node and two leaves, defined through a cut-point value for a single explanatory variable. In this way, we focus on uncovering just the main, most discriminant antibody for separating genotypes exhibiting a different phenotypical performance (see Fig. 4, SI *Appendix*, Fig. S8-S10). Once this was done, we focused on estimating the actual accuracy of the obtained model by conducting a 10-fold cross-validation process, i.e. the available data is randomly divided in 10 similar-size parts, and then similar trees are trained using 9 of these parts, leaving the remaining part to be used as a test sample (as these instances are not used in the tree fitting step). This step is carried out 10 times, each time using a different part as test sample. The proportion of test-sample instances correctly classified throughout this 10-steps process provides an estimation of the original model's accuracy. To obtain a more robust estimation and average out the dependence on the random division step, the whole 10-fold cross-validation process was replicated 100 times, each time using a different random division of the available data in 10 parts. The mean and standard deviation of these 100 10-fold cross-validation estimations at each addressed classification task are reported in Figure SI8B and Table S2. Both the CRT models fit and their cross-validation were performed using Python *scikit-learn* library.

***Glycome anlysis by Enzyme-Linked ImmunoSorbent Assay (ELISA)***

mAbs were obtained as hybridoma cell culture supernatants either from laboratory stocks (CCRC series, JIM series, MAC series; available from CarboSource [http://www.carbosource/net]) or from Plant Probes (LM series, PAM1 [http://www.plantprobes.net/]) unless otherwise indicated. A detailed list of all mAbs included in this study showing the immunogens used to develop them, their isotype, and the cell wall polysaccharide class they primarily recognize is provided in *Appendix*, Table S1. The experimental protocol previously described (2) was performed with minor modifications. In brief, flat-bottom 96-well ELISA plates (Nunc 439454 from Thermo Fisher Scientific or 2507 Costar from Corning Life Sciences), we used to apply cell wall mutant fraction polysaccharides (50 µL of 10 µg mL−1 in deionized water per well, or deionized water for controls) and then plates were dried to the well surfaces by evaporation overnight at 37°C. The plates were blocked with 200 µL of 1% (w/v) instant nonfat dry milk (Carnation) in Tris-buffered saline (50 mm Tris-HCl, pH 7.6, containing 100 mm sodium chloride) for 1 h. All subsequent aspiration and wash steps were performed using an ELx405 microplate washer (Bio-Tek Instruments). Blocking agent was removed by aspiration, and 50 µL of undiluted hybridoma supernatant of each antibody were added to the well and incubated for 1 h at room temperature. Supernatant was removed and wells were washed three times with 300 µL of 0.1% (w/v) instant nonfat dry milk in Tris-buffered saline (wash buffer). Peroxidase-conjugated goat anti-mouse IgG or goat anti-rat IgG antibodies (Sigma-Aldrich), depending on the primary antibody used, was diluted 1:5,000 in wash buffer, and 50 µL were added to each well and incubated for 1 h. Note that the secondary antibodies used in this study are generated against whole immunoglobin molecules and thus bind to several isotypes of primary antibodies, including IgGs, IgMs, and IgAs, according to the manufacturers. Wells were then washed five times with 300 µL of wash buffer. 3,3′,5,5′-Tetramethylbenzidine substrate solution (Vector Laboratories) was freshly prepared according to the manufacturer's instructions, and 50 µL were added to each well. After 20 min, the reaction was stopped by adding 50 µL of 0.5 n sulfuric acid to each well. The OD of each well was read as the difference in A450 and A655 using a model 680 microplate reader (Bio-Rad). The reading from each test well was subtracted from that of a control well on the same plate that contained the same primary and secondary antibodies but no immobilized polysaccharide. This experiment was repeated 3 times with 2 independent biological replicates.

| Locus[1] | Allele Tested[2] | Allele Mutant Line[3] | Ecotype | Protein Function | Reference[4] |
|---|---|---|---|---|---|
| **At1g12840** | **det3-1** | T==> A in 1[er] intron | Col-0 | Subunit C vacuolar H+ATPase (V-ATPase)/ De-ETiolated 3 (DET3) | Luo *et al*., 2015 (3) |
| | det3-2 | SAIL_517_E02 | | | Delgado-Cerezo *et al*., 2012 (4) |
| **At1g20850** | **xcp2-1** | SALK_057921.45.15.x | Col-0 | Papain type cystein endopeptidase/ Xylem cysteine protease 2 (XCP2) | Funk *et al*., 2002 (5) |
| | xcp2-2 | SALK_010938.56.00.x | | | Zhang *et al*., 2014(6) |
| **At1g23170** | at1g23170-1 | SALK_046821.54.50.x | Col-0 | Protein of unknown function DUF2359; Transmembrane protein | Coates, 2003 (7) |
| | at1g23170-2 | SALK_056368.55.50.x | | | |
| **At1g27440** | **irx10-1** | SALK_055673.55.00.x | Col-0 | Glycosyl transferase family 47 | Persson et al., 2005 (8) Brown et al., 2005 (9) |
| At1g56330 | sar1b-4 | Versailles_141E05 | Ws-0 | Sec23a/Small GTP-binding protein/ ARF-like GTPase family (SAR1b) | Zeng et al., 2015 (10) |
| At1g56340 | crt1-2 | Versailles_048A05 | Ws-0 | Calreticulin 1 (CRT1) | Christensen et al. 2010 (11) |
| **At1g65580** | **fra3-1** | A833V | Col-0 | Type II inositol polyphosphate 5-phosphatases/ FRAgile Fibre 3 (FRA3) | Zhong et al., 2004 (12) |
| **At1g69530** | **exp1-1** | Versailles_401A10 | Ws-0 | Expansin 1 (At-EXP1) | Esmon et al., 2005 (13) |
| **At1g70770** | **at1g70770-1** | SALK_023673.43.50.x | Col-0 | Protein of unknown function DUF2359, Transmembrane Protein | Schmidt et al., 2007 (14) |
| | at1g70770-2 | SAIL_1186_D04 | | | |
| At1g73440 | at1g73440-1 | Versailles_302D07 | Ws-0 | Calmodulin-like protein | Day et al., 2002(15) |
| At1g75500 | wat1-1 | SALK_001389 | Col-0 | Walls Are Thin 1(WAT1) /Nodulin21 | Ranocha et al.,2010 (16) Denancé et al., 2013(17) |
| At2g02120 | pdf2.1-2 | SAIL_67_F04 | Col-0 | Plant defensin protein, PDF2.1 | Siddique et al., 2011 (18) |
| **At2g27040** | ago4-1t | SALK_007523.54.75.x | Col-0 | ARGONAUTE 4 (AGO4) | Agorio and Vera, 2007(19 |
| At2g36170 | ubq2-1 | Versailles_323G03 | Ws-0 | Ribosomal Protein L40A, RPL40A/ Ubiquitin extension protein 2 (UBQ2) | Kim et al., 2013(20) |
| At2g38080 | irx12-1 | SALK_051892 | Col-0 | Laccase 4/ IRregular Xylem 12 (IRX12) | Brown et al., 2005 (9) Yi Chou et al., 2018 (21) |
| At2g46030 | ubc6-1 | Versailles_236D07 | Ws-0 | Ubiquitin-conjugating enzyme 6 (UBC6) | Watts et al., 1994 (22) |
| At3g10740 | araf1-1 | FLAG_091G07 | Ws-0 | α-L-ARAbinoFuranosidase 1(ARAF1/ASD1) | Montes et al., 2008 (23) |
| At3g16920 | ctl2-1 | SALK_055713 | Col-0 | ChiTinase-Like protein 2 (CTL2) | Sánchez-Rodríguez et al., 2012 (24) |
| At3g47510 | at3g47510-1 | SALK_030833 | Col-0 | Transmembrane protein | N.a. |
| At3g53210 | mtn21-1 | Versailles_258G08 | Ws-0 | Nodulin MtN21family protein | Busov et al., 2004 (25) |
| At3g54920 | pmr6-1 | G140D | Col-0 | Pectato Lyae Like/Powdery Mildew Resistance 6 (PMR6) | Vogel et al., 2002 (26) |
| **At4g02380** | **sag21-1** | SALK_099663.54.70.x | Col-0 | Senescence Associated Gene 21 (SAG21) | Salleh et at., 2012 (27) |
| **At4g15160** | at4g15160-1 | SALK_007014.56.00.x | Col-0 | Protease inhibitor/Lipid Transfer protein (LTP) | N.a. |
| **At4g18780** | **irx1-6** | W114Stop | Col-0 | Cellulose Synthase 8 (CESA8)/ IRregular Xylem 1 (IRX1) | Taylor et al., 2000 (28) Hernández-Blanco et al., 2007 (29) |
| At4g34460 | agb1-1 | G==> T splice 1[er] exon | Col-0 | Arabidopsis heterotrimeric G Protein Β subunit 1 (AGB1) | Delgado-Cerezo et al., 2012 (4) Llorente et al., 2005(30) |
| **At4g37770** | **acs8-2** | SALK_066725.31.85.x | Col-0 | 1-aminocyclopropane-1-carboxylate synthase-like protein 8(ACS8) | Zhang et al., 2018(31) |
| | acs8-3 | SAIL_102_E05 | | | |
| **At5g04370** | namt1-1 | SALK_001690.56.00.x | Col-0 | Nicotinate MethylTransferase 1 (NaMT1) | Wu et al., 2018 (32) |
| **At5g15630** | **irx6-1** | FLAG_248_B03 | Ws-0 | COBRA-LIKE 4 (COBL4)/ IRregular Xylem 6 (IRX6) | Brown et al., 2005 (9) Delgado-Cerezo et al., 2012 (4) |
| **At5g17420** | **irx3-1** | SRT 206 | La-*er* | Cellulose Synthase 7(CESA7/MUR10) IRregular Xylem 3 (IRX3) | Turner & Somerville, 1997 (33) Hernández-Blanco et al., 2007 (29) |
| At5g18650 | miel1-1 | SALK_097638 | Col-0 | E3 ubiquitin-protein ligase MIEL1 | Marino et al., 2013 (34) |
| At5g26120 | araf2-1 | SALK_033343 | Col-0 | α-L-ARAbinoFuranosidase 2(ARAF1/ASD2) | Fulton & Cobbett, 2003 (35) |
| **At5g49720** | **irx2-1** | P250L | La-*er* | KORRIGAN1 (KOR1), IRregular Xylem 2, β,1-4 endoglucanase | Szyjanowicz et al., 2004 (36) López-Cruz et al., 2014 (37) |
| **At5g51890** | **at5g51890-1** | SALK_086448.49.80.x | Col-0 | Peroxidase/Orrtholog of ZPO-C | Sato et al., 2006 (38) |
| At5g54690 | irx8-1 | SALK_014026 | Col-0 | Galacturonosyltransferase (GAUT1) / IRregular Xylem 8 (IRX8) | Persson et al., 2007 (39) |
| At5g58600 | pmr5-1 | W265Stop | Col-0 | Pectin acetyltransferase/Powdery Mildew Resistance 5 (PMR5) | Vogel et al., 2005 (40) Chiniquy et al., 2019(41) |
| At5g60340 | akk6-1 | SALK_015289 | Col-0 | Adenylate Kinase isoenzyme 6 homolog (AAK6) | Slovak et al., 2020 (42) |
| **At5g62920** | arr6-2 | SALK_008866.55.00.x | Col-0 | Arabidopsis Response Regulator 6 (ARR6) | Bacete et al., 2020(43) |
| | **arr6-3** | SALK_133123.21.20.x | | | |
| At5g63670 | spt4-1 | Versailles_460C02 | Ws-0 | Transcription elongation factor SPT4 homolog 2 | Dürr et al., 2014(44) |

[1]In bold are indicated the genes whose mutants have been analyzed for fitness parameters (Fig. 2)

[2]In bold are indicated mutant alleles analyzed for fitness parameters (biomass and seed yield; Fig. 2); additional mutant alleles tested in the disease resistance against *PcBMM* are underlined (SI5)

[3]Reference of the mutant lines used is indicated (T-DNA insertion lines and point mutations in EMS-derived mutants, with amino acids or exon splicing changes)

[4]References describing a function of the indicated gene/protein in cell wall biogenesis/composition is indicated in green, in disease resistance in red and in both phenotypes in blue.

**Figure S1. *Arabidopsis thaliana m*utants used in the disease resistance analyses.**

| Locus | Mutant allele | *P. cucumerina* | | *H. arabidopsidis* | | *R. pseudosolanacearum* | |
|---|---|---|---|---|---|---|---|
| | | Disease Rating (0-5) | +/- SD | Conidiospores/mg fresh weight | +/- SD | Disease Rating (0-4) | +/- SD |
| *Ecotype* | Col-0 | 2.1 | 0.06 | 576.74 | 30.57 | 2.27 | 0.03 |
| ***Ecotype*** | **Ws-0** | 2.07 | 0.11 | 1661.64 | 122.65 | 2.17 | 0.06 |
| ***Ecotype*** | **La-er** | 2.07 | 0.12 | 969.31 | 172.52 | 3.02 | 0.23 |
| *At1g12840* | *det3-1* | 1.77 | 0.13 | 227.74 | 38.24 | 0.01 | 0.15 |
| *At1g20850* | *xcp2-1* | 2.21 | 0.13 | 0 | 63.18 | 1.48 | 0.15 |
| *At1g23170* | *at1g23170-1* | 2.26 | 0.12 | 278.09 | 63.18 | 2.45 | 0.15 |
| *At1g27440* | *irx10-1* | 2.06 | 0.16 | 508.09 | 38.24 | 0.18 | 0.19 |
| ***At1g56330*** | ***sara1b-4*** | 2.43 | 0.1 | 1280.9 | 186.8 | 2.31 | 0.3 |
| ***At1g56340*** | ***crt1-1*** | 2.67 | 0.23 | 1455.16 | 396.45 | 2.01 | 0.23 |
| *At1g65580* | *fra3-1* | 0.95 | 0.19 | 258.59 | 38.43 | 0.08 | 0.15 |
| ***At1g69530*** | ***exp1-1*** | 1.14 | 0.26 | 1849.42 | 122.93 | 2.38 | 0.12 |
| *At1g70770* | *at1g70770-1* | 1.57 | 0.15 | 361.43 | 63.18 | 2.25 | 0.15 |
| ***At1g73440*** | ***at1g73440-1*** | 1.54 | 0.31 | 5245.89 | 1253.9 | 2.09 | 0.19 |
| *At1g75500* | *wat1-1* | 1.05 | 0.19 | 1418 | 202.8 | 1.01 | 0.02 |
| ***At2g02120*** | ***pdf2.1-2*** | 2.28 | 0.16 | 1782.9 | 203.19 | 3.41 | 0.25 |
| *At2g27040* | *ago4-1t* | 1.6 | 0.16 | 550.84 | 122.93 | 2.65 | 0.13 |
| ***At2g36170*** | ***ubq2-1*** | 2.43 | 0.1 | 1451.3 | 268.7 | 2.08 | 0.2 |
| *At2g38080* | *irx12-1* | 2.2 | 0.15 | 611.48 | 60.2 | 2.3 | 0.18 |
| ***At2g46030*** | ***ubc6-1*** | 2.19 | 0.2 | 2136.86 | 472.56 | 2.23 | 0.25 |
| ***At3g10740*** | ***araf1-1*** | 2.17 | 0.18 | 1438.77 | 170.44 | 2.15 | 0.16 |
| *At3g16920* | *ctl2-1* | 1.9 | 0.15 | 436.84 | 72.69 | 1.85 | 0.17 |
| *At3g47510* | *at3g47510-1* | 1.51 | 0.12 | 976.15 | 79.8 | 2.2 | 0.12 |
| ***At3g53210*** | ***mtn21-1*** | 1.91 | 0.15 | 1385.23 | 152.23 | 1.98 | 0.25 |
| *At3g54920* | *pmr6-1* | 1.96 | 0.15 | 496.07 | 72.69 | 2.05 | 0.2 |
| *At4g02380* | *sag21-1* | 1.76 | 0.18 | 201.29 | 63.18 | 2.74 | 0.15 |
| *At4g15160* | *at4g15160-1* | 2.07 | 0.13 | 0 | 63.18 | 2.37 | 0.15 |
| *At4g18780* | *irx1-6* | 1 | 0.1 | 477.75 | 38.24 | 0.05 | 0.13 |
| *At4g34460* | *agb1-1* | 4.8 | 0.2 | 679.11 | 72.69 | 2.15 | 0.15 |
| *At4g37770* | *acs8-2* | 1.93 | 0.18 | 304.09 | 63.18 | 1.18 | 0.15 |
| *At5g04370* | *namt1-1* | 1.64 | 0.13 | 395.48 | 63.18 | 2.23 | 0.15 |
| ***At5g15630*** | ***irx6-1*** | 1.24 | 0.16 | 2059.12 | 123.78 | 2.02 | 0.16 |
| ***At5g17420*** | ***irx3-1*** | 1.08 | 0.18 | 845.76 | 173.52 | 0.35 | 0.08 |
| *At5g18650* | *miel1-1* | 2.23 | 0.15 | 883.67 | 344.5 | 2.63 | 0.12 |
| *At5g26120* | *araf2-1* | 2.36 | 0.14 | 1509.72 | 301.8 | 2.08 | 0.24 |
| ***At5g49720*** | ***irx2-1*** | 1.84 | 0.16 | 951.39 | 172.52 | 3 | 0.23 |
| *At5g51890* | *at5g51890-1* | 1.61 | 0.17 | 430.73 | 30.57 | 2.35 | 0.15 |
| *At5g54690* | *irx8-1* | 1.97 | 0.18 | 441.33 | 72.69 | 2.47 | 0.18 |
| *At5g58600* | *pmr5-1* | 1.9 | 0.2 | 615.88 | 72.69 | 2.3 | 0.21 |
| *At5g60340* | *akk6-1* | 2.42 | 0.15 | 1668.48 | 222.5 | 2.19 | 0.23 |
| *At5g62920* | *arr6-3* | 1.81 | 0.1 | 401.26 | 38.24 | 3.23 | 0.2 |
| ***At5g63670*** | ***spt4-1*** | 0.61 | 0.1 | 3815.47 | 852.26 | 1.95 | 0.28 |

Susceptible Wild type Resistant

**Figure S2. Disease resistance analysis of Arabidopsis cell wall mutants**. Average disease resistance values (+/- SD) of wild-type plants and cell wall mutants in different backgrounds (black-Col-0; green-La-*er*; purple-Ws-0) to *P. cucumerina* BMM, *R. pseudosolanacearum* (GMI1000 strain for Col-0 and La-*er*, and RD15 for Ws-0), and *H. arabidopsidis* (Noco2 strain for Col-0, Ewma1 for La-*er* and Cala for Ws-0). Color-code of the corresponding columns indicates the level of the resistance phenotype, from susceptible (blue) to resistant (red) in comparison to wild-type (white).

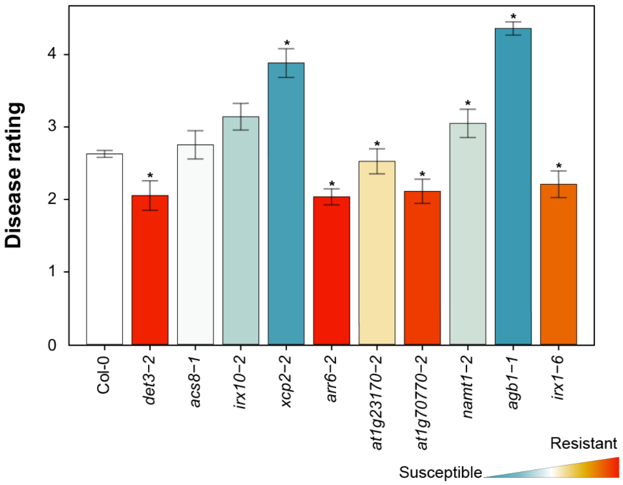


**Figure S3. Disease rating (DR) of *cwm* second alleles inoculated with *P. cucumerina* BMM**. DR average (±SD) of wild-type (wt) plants (Col-0 and Ws-0 backgrounds) and mutants at 7 days post inoculation (dpi) with the necrotrophic fungus *P. cucumerina* BMM. DR varies from 0 (non-infected plants) to 5 (dead plants). Asterisk indicate significant differences compared with wt values (ANOVA non-balanced analysis. Dunnet test p≤0.05). This is one representative experiment of the three performed that gave similar results (n = 10).

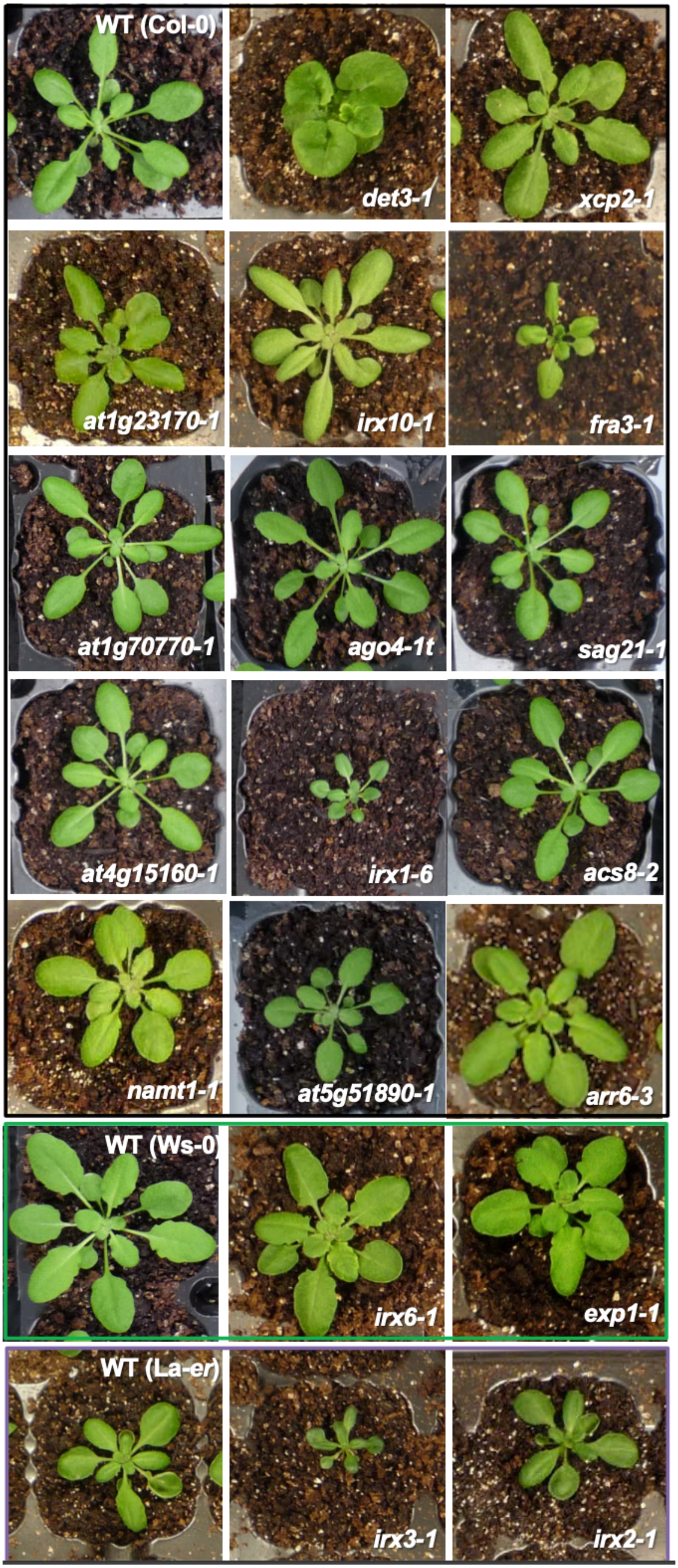


**Figure S4. Developmental phenotypes of three-week old plants of *cwm* and wild-type ecotypes (Col-0, Ws-0 and La-*er*).** Plants were grown under short day conditions as described in Material and Methods.

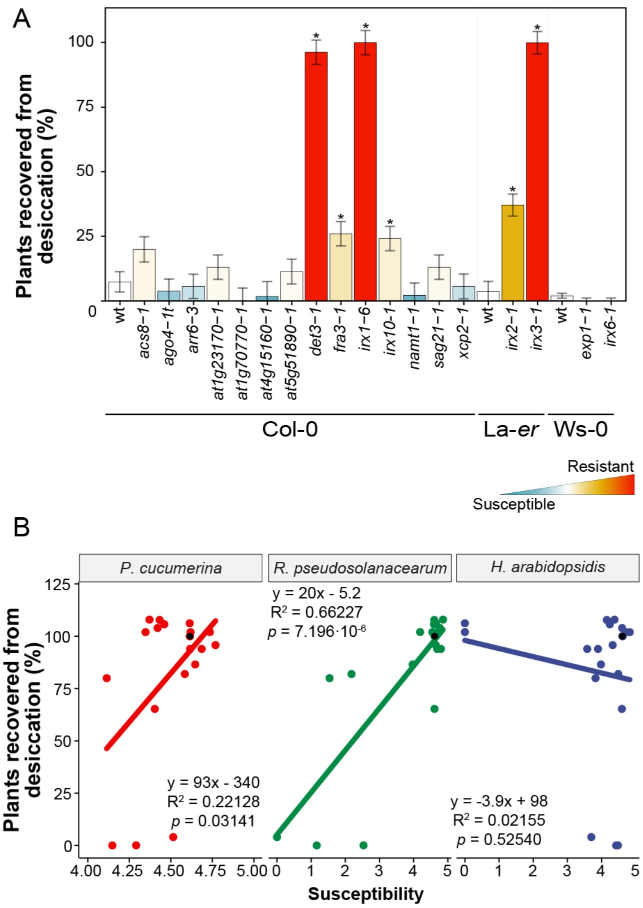


**Figure S5. Desiccation tolerance of wild-type plants and cell wall mutants.** *(A)* Percentage of plant survival after drought stress application for 21 days followed by plant re-watering. *(B)* Correlation analysis between desiccation tolerance and resistance to pathogens of 18 *cwm* mutants and wild-type plants genotypes. Average response information of each genotype (dot in the graph) is expressed in relation to that of the reference wild-type plant (black dot, value of 100% at the *y*-axes). Plants biotic stress susceptibility ratios were log-transformed, and accordingly *x*-axes range from 0 (lower susceptibility) to 5 (greater susceptibility), with the wild-type plants situated at 4.72 = *ln* (1 + 100). A linear model was fitted for each combination and correlations determined. Fitted equations, R-squares and *p*-values are indicated in the insets of graphs.

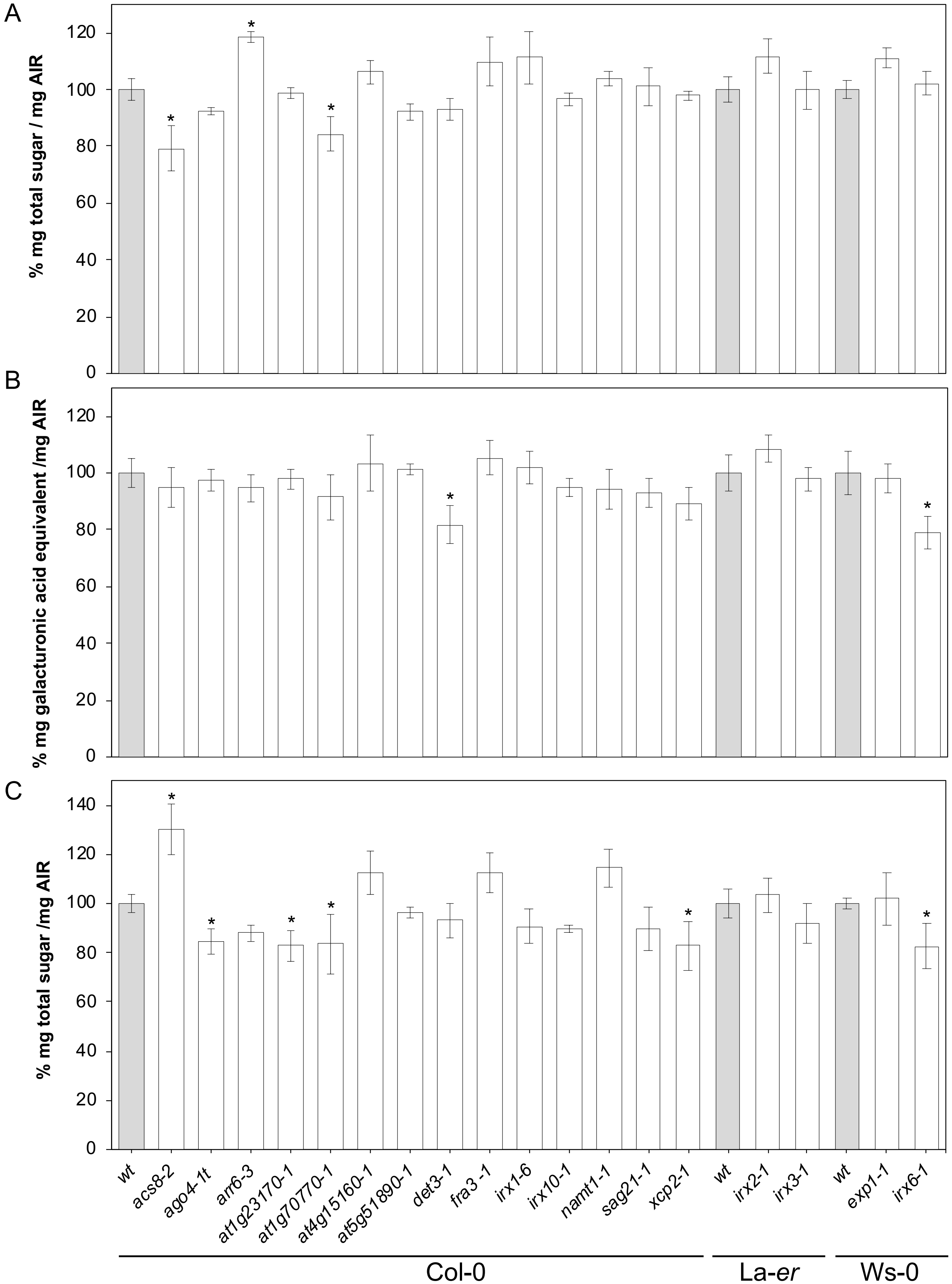


**Figure S6. Cell wall biochemical composition of *Arabidopsis cwm* plants.** (*A*) Total sugars quantification (% µg per mg of dry weight) in the non-cellulosic carbohydrate fraction from the cell walls of the mutants and their corresponding background plants (Col-0, La-*er* and Ws-0). (*B*) Total uronic acid (UA) content (% µg per mg of dry weight) from the cell walls of the indicated genotypes. (*C*) Cellulose content (% µg of total sugars per mg of dry weight from the cell walls of the indicated genotypes). Data represent average values (± SE) of three independent experiments. Asterisks indicate mean values significantly different from wild-type plants (Student's *t*-test. $p < 0.1$, n > 10).

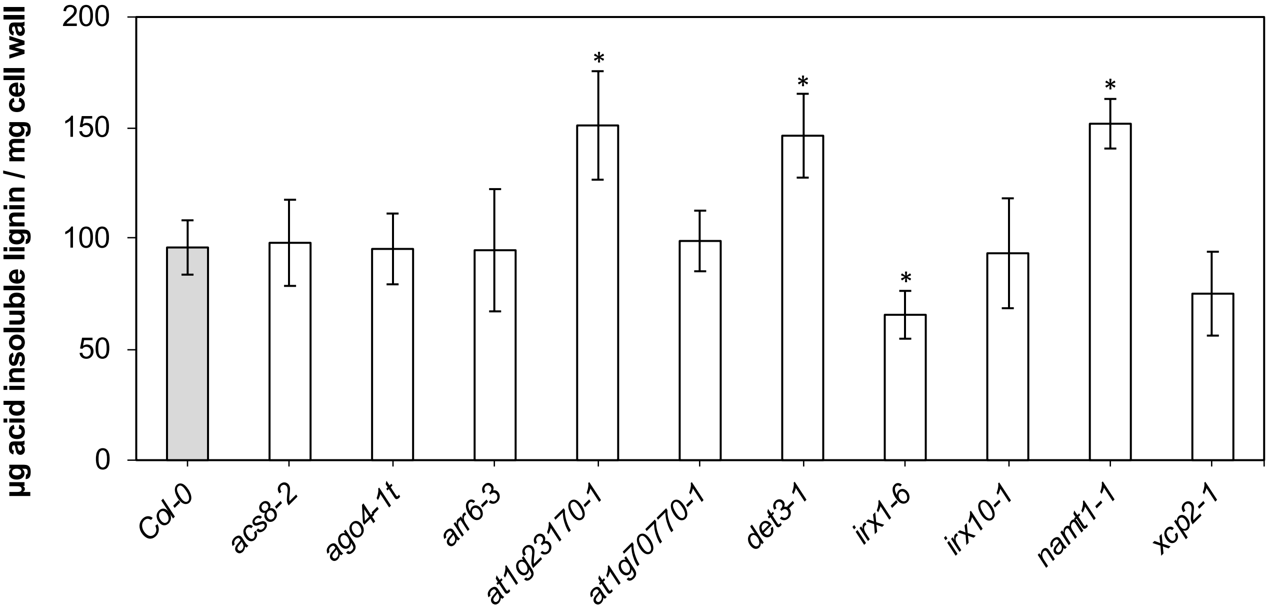


**Figure S7. Total lignin content of *Arabidopsis cwm* plants**. Quantification of total lignin (µg acid insoluble lignin per mg cell wall) in the indicated mutants and their corresponding background plant (Col-0). Data represent average values (± SE) of three independent experiments. Asterisks indicate mean values significantly different from wild-type plants (Student's *t*-test. $p < 0.1$, n > 10).

A

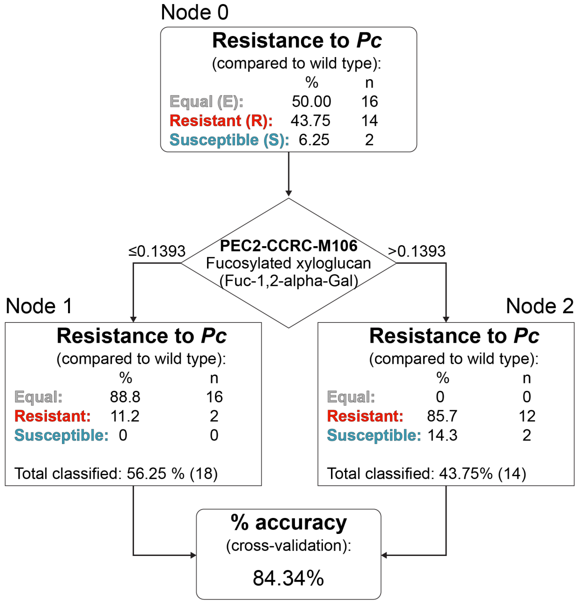


B

| Phenotype | CW Fraction | Antibody | Abs. cut - off | Classification (compared to wt) R/H | E | S/L | Accuracy |
|---|---|---|---|---|---|---|---|
| Resistance to *P. cucumerina* | PEC2 | CCRC_M106 | ≤ 0.1393 | 2 | 16 | 0 | 84.34% |
| | | | > 0.1393 | 12 | 0 | 2 | |
| Resistance to *R. pseudosolanacearum* | PEC2 | CCRC_M5 | ≤ 0.2990 | 10 | 0 | 2 | 83.88% |
| | | | > 0.2990 | 0 | 18 | 2 | |
| Resistance to *H. arabidopsidis* | PEC2 | CCRC_M106 | ≤ 0.1300 | 4 | 12 | 0 | 83.43% |
| | | | > 0.1300 | 16 | 0 | 0 | |
| | HEC1 | CCRC_M174 | ≤ 0.0005 | 0 | 7 | 0 | 82.96% |
| | | | > 0.0005 | 20 | 5 | 0 | |
| Rosette biomass | PNS | CCRC_M22 | ≤ 0.4195 | 0 | 13 | 0 | 87.31% |
| | | | > 0.4195 | 0 | 3 | 16 | |
| Seed production | HEC1 | CCRC_M175 | ≤ 0.0052 | 0 | 21 | 0 | 87.62% |
| | | | > 0.0052 | 2 | 1 | 8 | |
| | PEC1 | CCRC_M170 | ≤ 0.0367 | 1 | 20 | 0 | 85.05% |
| | | | > 0.0367 | 1 | 2 | 8 | |
| Tolerance to desiccation | HEC2 | JIM 101 | ≤ 0.0040 | 2 | 24 | 0 | 83.16% |
| | | | > 0.0040 | 6 | 0 | 0 | |

**Figure S8. Predictive CRT model correlating wall composition and disease resistance/fitness/desiccation phenotypes of *Arabidopsis* cell wall mutants**. (*A*) Scheme of the CRT model obtained in the analysis of the correlation between resistance to *P. cucumerina* and cell wall epitopes. The tree model obtained with antibody CCRC-M106 (fucosylated xyloglucan) in PEC2 fraction of *cwm* and wild-type plants is shown. (*B*) Summary of the most relevant CRT models obtained for the different variables (resistance to pathogens, fitness and desiccation tolerance). For each variable, the CRT-selected antibodies detecting epitopes of some cell wall extracts as well as their absorbance cut-points are indicated (see A). The number of observations (*cwm* and wild-type plants) of each disease resistance/fitness/desiccation phenotype (resistant/higher (R/H), equal (E) or susceptible/lower (S/L) than wild-type plants) verifying either side of the cut-points is also reported. The predictive accuracy (% accuracy) of each model is estimated as the percentage of correct classifications

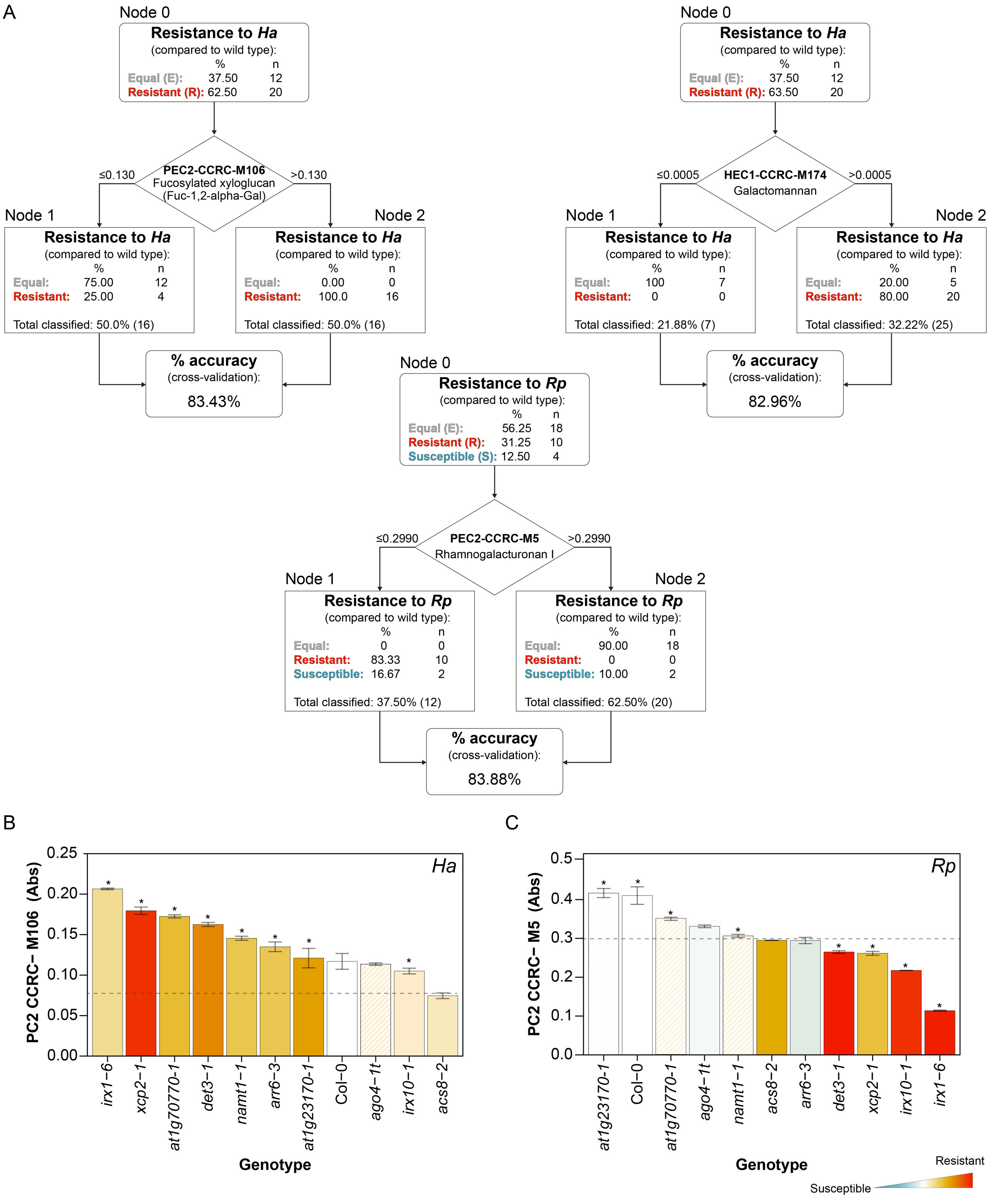


**Figure S9. Predictive CRT model correlating wall composition and disease resistance/fitness phenotypes of *Arabidopsis* cell wall mutants to *H. arabidopsidis and R. pseudosolanacearum*.** *(A)* Scheme of the CRT models obtained in the analysis of the correlation between resistance to *H. arabidopsidis and R. pseudosolanacearum* and cell wall epitopes. The tree models obtained with antibodies in cell wall fractions of *cwm* and wild-type plants are shown. *(B)* Biological validation of CRT results with cell wall mutants from 6 clusters analyzed. The absolute value (average ±SD) of the epitope signal detected by the antibody are shown (*n* = 3). The color code of the column indicates the resistance level of the corresponding mutant, from red (resistant) to blue (susceptible) in comparison with wild-type (wt) resistance (in white). Stripped columns mean no significant resistance differences with wt plants (see Fig. 1). The absorbance cut-point value for considering a mutant as resistant as determined by CRT is indicated by the dotted lines. The *cwm* lines that fulfill the CRT model are marked with an asterisk.

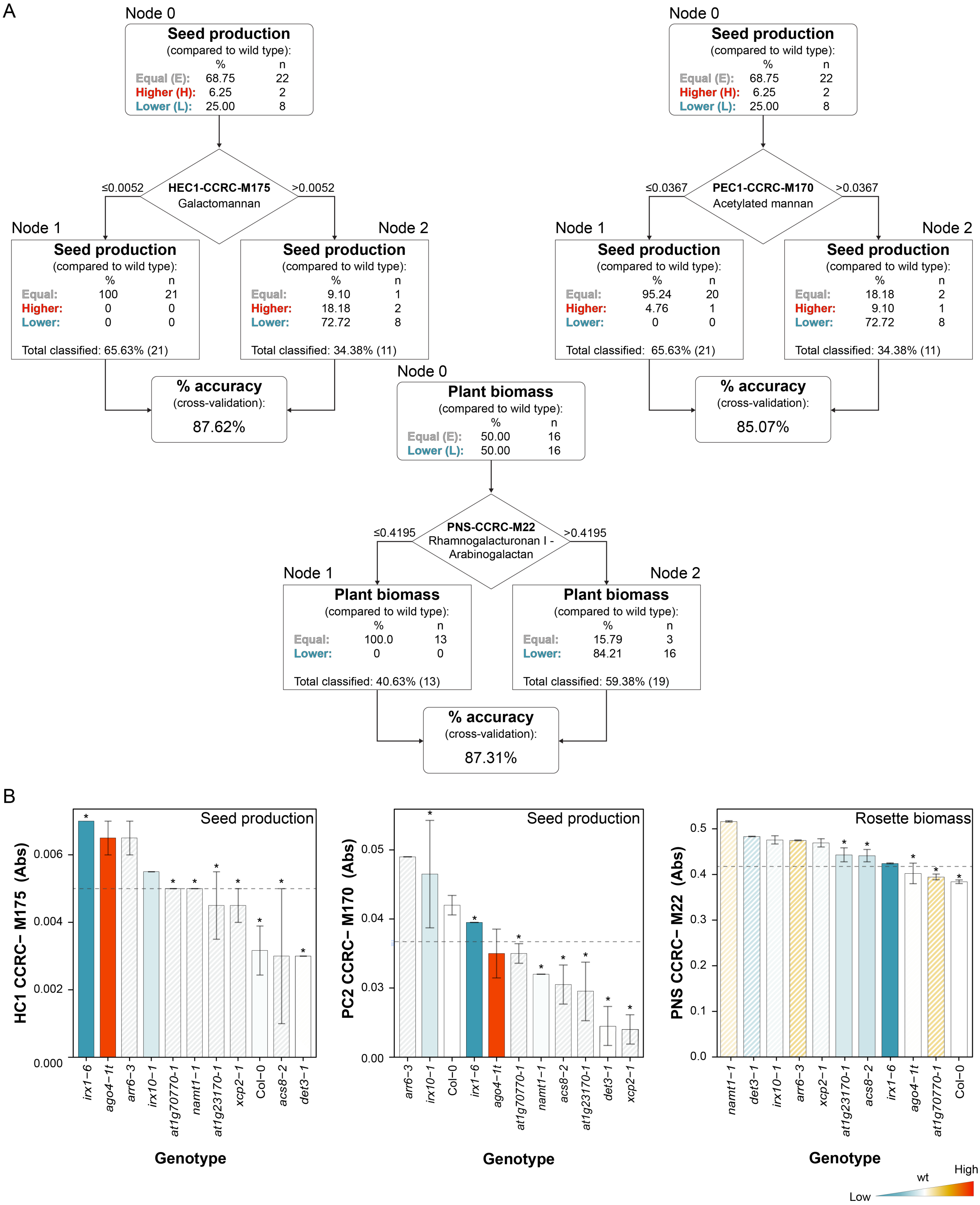


**Figure S10. Predictive CRT model correlating wall composition and fitness phenotypes of *Arabidopsis* cell wall mutants**. *(A)* Predictive CRT model correlating wall composition and fitness phenotypes of *Arabidopsis* cell wall mutants. *(A)* Scheme of CRT models obtained in the analysis of the correlation between fitness and cell wall epitopes. The tree models obtained with antibodies in cell wall fractions of *cwm* and wild-type plants are shown. *(B)* Biological validation of CRT results with cell wall mutants analyzed*.* The absolute value (average ±SD) of the epitope signal detected by the antibody are shown (n = 3). The color code of the column indicates the level of the fitness value corresponding to mutant, from red (enhanced values) to blue (reduced values) in comparison with wild-type (wt) resistance (in white). Stripped columns mean no significant fitness differences with wt plants (see Fig. 2A,B) The absorbance cut-point value for considering a mutant as matching the CRT model is indicated by the dotted lines. The *cwm* lines that fulfill the CRT model are marked with an asterisk.

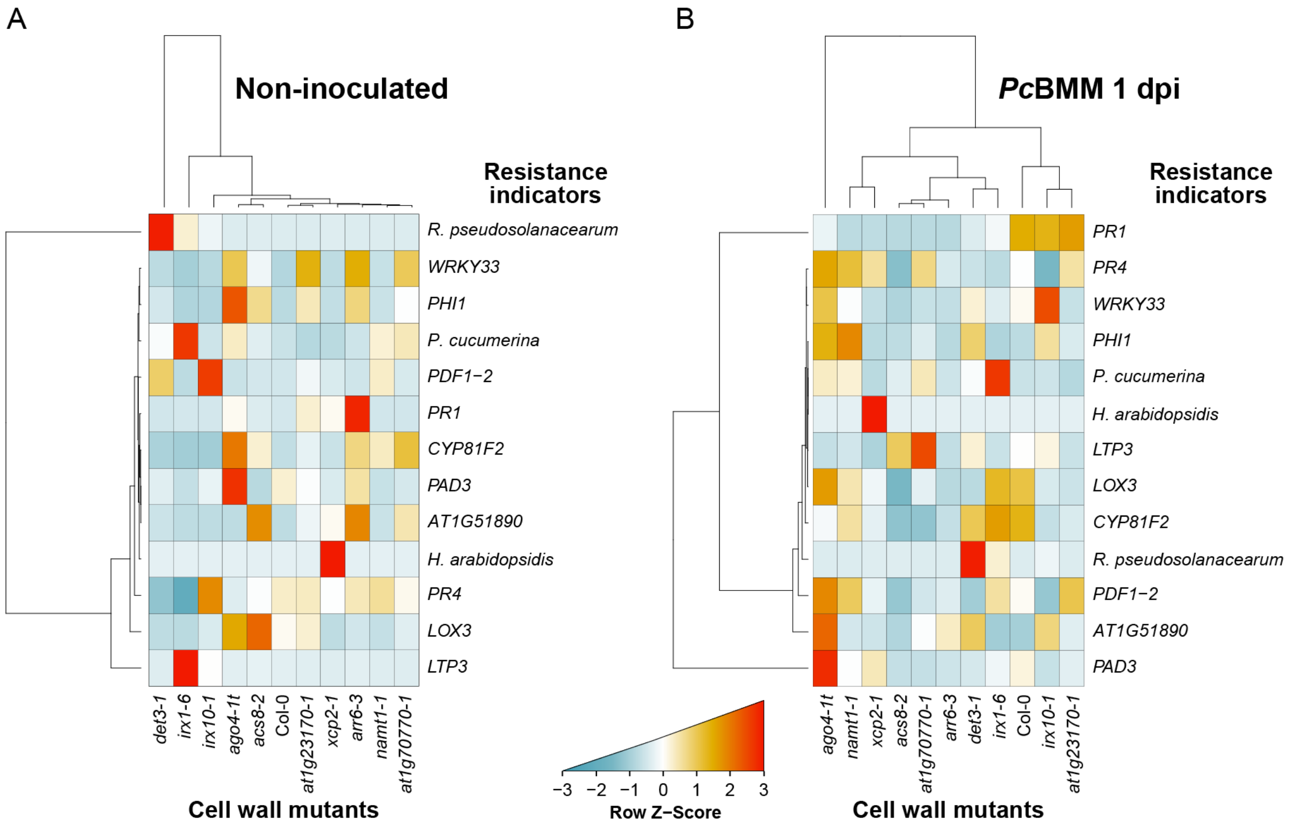


**Figure S11. Clustering of the expression pattern of canonical defensive genes in wild-type plants and *cwm* mutants and their disease resistance phenotypes**. (*A*) Clustering of disease resistance phenotypes and expression levels (determined by qRT-PCR) of defensive and MAMP-induced genes in non-inoculated three-week old plants from the indicated genotypes. (*B*) Relative expression (*PcBMM*/Mock 1dpi). Clusters were computed using Euclidean distances for absolute gene expression levels and disease indexes, and Z-scores were calculated across rows for normalization. Expression levels relative to the *UBC21* gene and mock-treatment are shown. Values are means ± SE (*n* = 3). Experiments were performed three times with similar results.

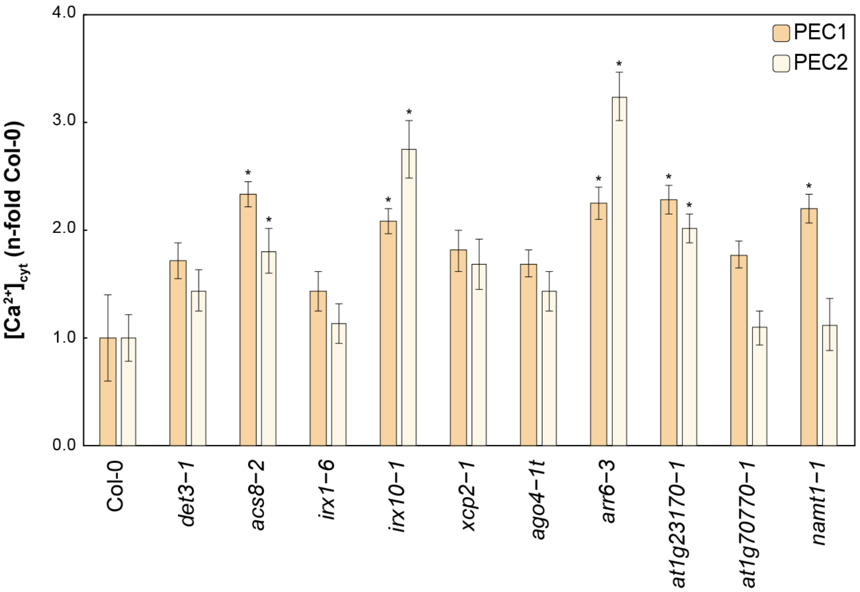


**Figure S12. Cell wall derived extracts from pectin fractions (PEC1 and PEC2) of *cwm* plants trigger $Ca^{2+}$ elevations.** Increases in cytoplasmic calcium concentrations ($[Ca^{2+}]_{cyt}$) in Col-0AEQ seedlings upon treatment with heat treated (121 ºC for 20 min) cell walls extracts (PEC1and PEC2) from Col-0 and the indicated cell wall mutants plants. PEC1: weakly bound pectin fraction (50 ng/µl). PEC2: highly bound pectin fraction (50 ng/µl). Data are representative of three independent experiments with similar results (means ± SD, n = 16). Asterisks indicate statistically significant differences ($p$ < 0.05) with the corresponding fraction from Col-0 plants (ANOVA, Dunnett's multiple comparisons test correction).

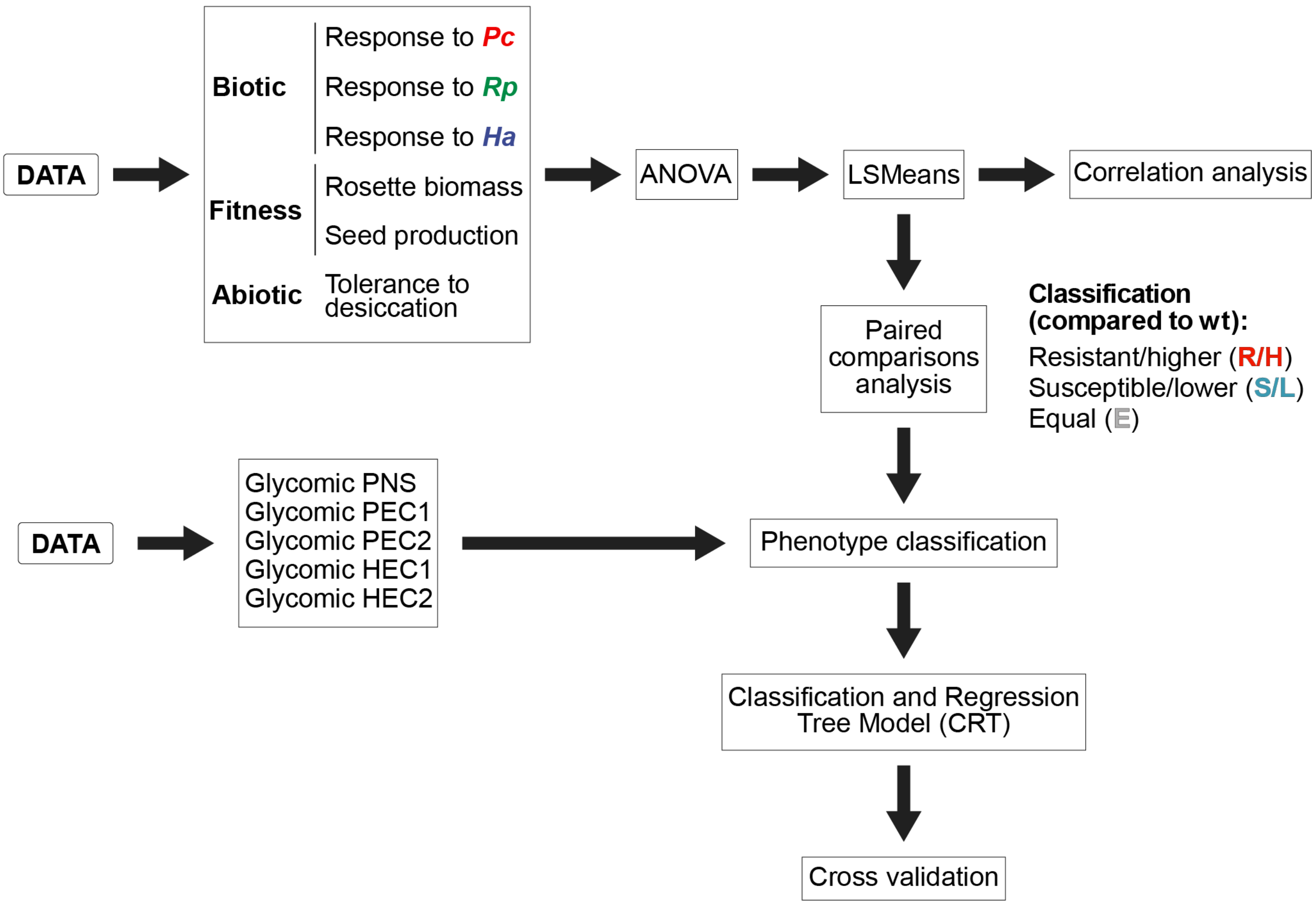


**Figure S13. Schema of the mathematical analyses performed to generate the data of Fig. 2C, Fig. 4, Fig. S5B, Fig S8-S10.**

**Table S2: Summary of the results from the performed Classification and Regression Tree (CRT) analysis.** The observed phenotypes and the used cell wall fractions are indicated. The monoclonal antibodies which provided a better classification of the observed phenotypes were selected, and the epitope they recognise is shown. Cut-off values are absorbance values that determine the two classification categories. Raw and corrected accuracy values are shown.

| Phenotype | Wall Fraction | Antibody | Recognised wall Epitope | Cut_point | Leaf1_ERS | Leaf2_ERS | OK_E/Total_E | OK_R/Total_R | OK_S/Total_S | Accuracy | CV_Accuracy | CV_AccStd |
|---|---|---|---|---|---|---|---|---|---|---|---|---|
| *Pc* | PNS | CCRC_M42 | Rhamnogalacturonan I | 0.1757 | 11/1/0 | 5/13/2 | 11/16 | 13/14 | 0/2 | 75 | 47.95 | 6.19 |
| *Pc* | PEC1 | CCRC_M30 | Rhamnogalacturonan I | 0.0865 | 0/11/0 | 16/3/2 | 16/16 | 11/14 | 0/2 | 84.375 | 78.53 | 2.60 |
| *Pc* | PEC2 | CCRC_M106 | Fucosylated xyloglucan (Fuc-1,2-alpha-Gal) | 0.1393 | 16/2/0 | 0/12/2 | 16/16 | 12/14 | 0/2 | 87.5 | 84.34 | 4.18 |
| *Pc* | HEC1 | CCRC_M174 | Galactomannan | 0.0022 | 15/2/1 | 1/12/1 | 15/16 | 12/14 | 0/2 | 84.375 | 76.35 | 4.71 |
| *Pc* | HEC2 | CCRC_M56 | Rhamnogalacturonan I | 0.6030 | 13/1/0 | 3/13/2 | 13/16 | 13/14 | 0/2 | 81.25 | 71.07 | 5.66 |
| *Rp* | PNS | CCRC_M25 | Rhamnogalacturonan I/Arabinogalactan | 0.2795 | 16/1/2 | 2/9/2 | 16/18 | 9/10 | 0/4 | 78.125 | 53.21 | 4.46 |
| *Rp* | PEC1 | JIM7 | Homogalacturonan (GalA1->4MeGalA1->4MeGalA1->4MeGalA1->4MeGalA1->4GalA) | 0.8295 | 17/3/4 | 1/7/0 | 17/18 | 7/10 | 0/4 | 75 | 49.74 | 5.25 |
| *Rp* | PEC2 | CCRC_M5 | Rhamnogalacturonan I | 0.2990 | 0/10/2 | 18/0/2 | 18/18 | 10/10 | 0/4 | 87.5 | 83.88 | 1.53 |
| *Rp* | HEC1 | CCRC_M114 | Xylan | 0.0118 | 0/0/4 | 18/10/0 | 18/18 | 0/10 | 4/4 | 68.75 | 51.97 | 6.61 |
| *Rp* | HEC2 | CCRC_M69 | Rhamnogalacturonan I | 0.2380 | 17/2/4 | 1/8/0 | 17/18 | 8/10 | 0/4 | 78.125 | 50.10 | 5.42 |
| *Ha* | PNS | JIM11 | Arabinogalactan | 0.0543 | 1/16/0 | 11/4/0 | 11/12 | 16/20 | 0/0 | 84.375 | 61.12 | 6.06 |
| *Ha* | PEC1 | CCRC_M26 | (1,3)(1,6)glucan | 0.0030 | 5/20/0 | 7/0/0 | 7/12 | 20/20 | 0/0 | 84.375 | 72.01 | 5.45 |
| *Ha* | PEC2 | CCRC_M106 | Fucosylated xyloglucan (Fuc-1,2-alpha-Gal) | 0.1300 | 12/4/0 | 0/16/0 | 12/12 | 16/20 | 0/0 | 87.5 | 83.43 | 2.43 |
| *Ha* | HEC1 | CCRC_M174 | Galactomannan | 0.0005 | 7/0/0 | 5/20/0 | 7/12 | 20/20 | 0/0 | 84.375 | 82.96 | 3.62 |
| *Ha* | HEC2 | CCRC_M138 | Xylan | 0.2002 | 4/20/0 | 8/0/0 | 8/12 | 20/20 | 0/0 | 87.5 | 81.34 | 5.75 |
| Biomass | PNS | CCRC_M22 | De-arabinosylated rhamnogalacturonan I | 0.4195 | 13/0/0 | 3/0/16 | 13/16 | 0/0 | 16/16 | 90.625 | 87.31 | 1.56 |
| Biomass | PEC1 | CCRC_M117 | Xylan | 0.0012 | 10/0/1 | 6/0/15 | 10/16 | 0/0 | 15/16 | 78.125 | 45.32 | 5.54 |
| Biomass | PEC2 | CCRC_M5 | Rhamnogalacturonan I | 0.3185 | 2/0/12 | 14/0/4 | 14/16 | 0/0 | 12/16 | 81.25 | 52.67 | 4.92 |
| Biomass | HEC1 | CCRC_M154 | Xylan | 0.0075 | 10/0/0 | 6/0/16 | 10/16 | 0/0 | 16/16 | 81.25 | 64.38 | 3.89 |
| Biomass | HEC2 | CCRC_M69 | Rhamnogalacturonan I | 0.2072 | 11/0/1 | 5/0/15 | 11/16 | 0/0 | 15/16 | 81.25 | 54.39 | 6.05 |
| Seeds | PNS | CCRC_M128 | Rhamnogalacturonan I | 0.2570 | 20/0/2 | 2/2/6 | 20/22 | 0/2 | 6/8 | 81.25 | 54.73 | 6.42 |
| Seeds | PEC1 | CCRC_M170 | Acetylated mannan | 0.0367 | 20/1/0 | 2/1/8 | 20/22 | 0/2 | 8/8 | 87.5 | 85.07 | 3.71 |
| Seeds | PEC2 | CCRC_M106 | Fucosylated xyloglucan (Fuc-1,2-alpha-Gal) | 0.1920 | 22/2/2 | 0/0/6 | 22/22 | 0/2 | 6/8 | 87.5 | 77.53 | 4.78 |
| Seeds | HEC1 | CCRC_M175 | Galactomannan | 0.0052 | 21/0/0 | 1/2/8 | 21/22 | 0/2 | 8/8 | 90.625 | 87.62 | 1.15 |
| Seeds | HEC2 | JIM19 | Arabinogalactan | 0.1258 | 22/1/2 | 0/1/6 | 22/22 | 0/2 | 6/8 | 87.5 | 62.39 | 5.07 |
| Drought | PNS | CCRC_M93 | Xyloglucan | 0.0052 | 2/6/0 | 22/2/0 | 22/24 | 6/8 | 0/0 | 87.5 | 58.43 | 6.08 |
| Drought | PEC1 | CCRC_M55 | Non-fucosylated xyloglucans. | 0.0155 | 17/0/0 | 7/8/0 | 17/24 | 8/8 | 0/0 | 78.125 | 51.74 | 6.41 |
| Drought | PEC2 | CCRC_M5 | Rhamnogalacturonan I | 0.2990 | 4/8/0 | 20/0/0 | 20/24 | 8/8 | 0/0 | 87.5 | 65.60 | 4.61 |
| Drought | HEC1 | CCRC_M7 | Rhamnogalacturonan I (trimer or larger of beta-(1,6)-Gal carrying one or more Ara residues of unknown linkage) | 0.4590 | 24/5/0 | 0/3/0 | 24/24 | 3/8 | 0/0 | 84.375 | 56.70 | 5.93 |
| Drought | HEC2 | JIM101 | Rhamnogalacturonan I | 0.0040 | 24/2/0 | 0/6/0 | 24/24 | 6/8 | 0/0 | 93.75 | 83.16 | 4.07 |

**Table S3: Oligonucleotides used for T-DNA insertional mutant characterization.**

| Gene | AGI locus | Background | Allele | Line | Forward oligonucleotide | Reverse oligonucleotide |
|---|---|---|---|---|---|---|
| *DET3* | *AT1G12840* | Col-0 | *det3-1* | CS6160 | Point mutation (T->A) in 1st intron 32 bp upstream of putative 3′ splice site | |
| | | | *det3-2* | SAIL_517_E02 | ATCCTCTTGCTCCTCTTCAGC | CTGCGAAATTGAAACCAAAAC |
| *XCP2* | *AT1G20850* | Col-0 | *xcp2-1* | SALK_057921.45.15.x | GACACTGAGAGGCTGATGAGC | AGCGACCTCTATCGAGTCTCC |
| | | | *xcp2-2* | SALK_010938.56.00.x | AAAGGGAAAAGCTACTGGCTC | GGTTTCCCAGTGTTCCTCTTC |
| *AT1G23170* | *AT1G23170* | Col-0 | *at1g23170-1* | SALK_046821.54.50.x | TTACGCAAAACCATTGCTACC | GGGAAGATCTAAATGGCGATC |
| | | | *at1g23170-2* | SALK_056368.55.50.x | ACCTTTTGCCTCAAGCTCTTC | ACAGTTTGGATGATGGCTCAG |
| *IRX10* | *AT1G27440* | Col-0 | *irx10-1* | SALK_055673.55.00.x | ACAAAAGCCGTGATCAATGAC | AAACATCACCAGCACTTCCTG |
| | | | *irx10-2* | SALK_046368.54.75.x | ACCATGTCTGTTTGGACGAAG | AAAATCCACTCGGAGGACTTG |
| *FRA3* | *AT1G65580* | Col-0 | *fra3-1* | SAIL_253_E02 | TTTGTAAATGAACGTCCCTGC | TACCAACAAATTCCGGTAACG |
| *EXP1* | *AT1G69530* | Ws-0 | *exp1-1* | Versalles_401A10 | | |
| | | | *exp1-2* | SALK_010506.54.50.x | CAAAGCAGACCACTATGACCC | TGTTCGGTAAGGCGTTGTTAG |
| *AT1G70770* | *AT1G70770* | Col-0 | *at1g70770-1* | SALK_023673.43.50.x | CAGGAAGTTCCCAAGAGATCC | CCGAATGATGCTCTCACTCTC |
| | | | *at1g70770-2* | SAIL_1186_D04 | GTCATCTGTCCGCAAAATGAG | ATCCTTTGGGATTTGGTTTTG |
| *AGO4* | *AT2G27040* | Col-0 | *ago4-1t* | Versalles_177B08 | | |
| | | Ws-0 | *ago4-3* | SALK_007523.54.75.x | TTTTGGTTCCAATTGCATCTG | CCCACAAAATCAAAGTGAAGAAG |
| *IRX9* | *AT2G37090* | Col-0 | *irx9-1* | SALK_058238.37.75.x | CCAAAACTGTCAATTTATAACATTGG | ATGTTCAATGTGCCTCAAAGC |
| | | | *irx9-2* | SALK_057033.51.30.x | GCTGGTAAGGCCTCATTTTTC | AACTTACCAACCCACCCATTC |
| *CTL2* | *AT3G16920* | Col-0 | *ctl2-1* | SALK_055713.38.85.x | CAGCTTCTTCTCGTCCAACAC | GTTTCGAAACCGCTATTCTCC |
| | | | *ctl2-2* | SAIL_545_A07 | TTGCTTAAGCCCAACAAAATG | AAGAAAATTGGTCGAAGCATG |
| *PMR6* | *AT3G54920* | Col-0 | *pmr6-1* | SAIL_519_C12 | TCCCATCTTTTCGACAAAATG | GTGTTGGTGAGGGAGAGTGAG |
| | | | *pmr6-2* | SALK_130726.44.10.x | TGAGCCAATGAATCAATCAATC | TGTGGAGGTTAAGGACGATTG |
| *SAG21* | *AT4G02380* | Col-0 | *sag21-1* | SALK_099663.54.70.x | TGGGTCAAAAGACTCAAAGGC | TATGCCAATCAAATTGGAACG |
| | | | *sag21-2* | SAIL_134_E11 | AGAGGATCAGGATTCAGGACAG | AATGATCTGGTGCAGTGAACC |
| *AT4G15160* | *AT4G15160* | Col-0 | *at4g15160-1* | SALK_007014.56.00.x | TAGTTTCCCAAATTTTTCGGG | GCCCTGGTCAAACAACTAGTG |
| | | | *at4g15160-2* | SALK_044936.30.00.x | TTTAAGCCCGGATAAACTTGG | TCCTCGAGTTTTTGTAAGGGAG |
| *IRX1* | *AT4G18780* | La-*er* | *irx1-1* | SALK_100960.55.50.x | CTGAGCAAAATCAGAAGGTCG | TCTTCACCACAAGTGTTGCAG |
| | | | *irx1-6* | SALK_046685.56.00.x | GAAATTTGTGTCGAGACCAGC | TACAGTCCACCTTCAAAACCG |
| *AGB1* | *AT4G34460* | Col-0 | *agb1-1* | | | |
| | | | *agb1-2* | | | |
| *ACS8* | *AT4G37770* | Col-0 | *acs8-1* | SALK_066725.31.85.x | ATAACCAACCCATCTAACCCG | GGCTTCTCAACCAGAAAGGTC |
| | | | *acs8-2* | SAIL_102_E05 | TCTTGTTCTTGTTCCCATTGG | TCTTCCCAACCCCAAAAATAC |
| *NAMT1* | *AT5G04370* | Col-0 | *namt1-1* | SALK_001690.56.00.x | GGCCAAGATCATATCCATGTG | TTTTGGTGCGATTTTGGATAC |
| | | | *namt1-2* | SAIL_300_D11 | CAATAGCCAGTACCACAACCC | AACAGAGGAACCAAAACCCAC |
| *IRX6* | *AT5G15630* | Ws-0 | *irx6-1* | SALK_096418.49.95.x | TGGAGGCAAATTTCAACAAAC | ATGTGCAATCAGAGGTTTTGC |
| *IRX3* | *AT5G17420* | La-*er* | *irx3-1* | SALK_029940.54.20.x | AGAGAAGCTTAAGGAAACCGC | GAACAACACAAGAGCAGAGGG |
| *IRX2* | *AT5G49720* | La-*er* | *irx2-1* | SALK_075812.55.50.x | TAGCTGCCCATATATTTTCGG | CAGTCCAGACGAAGATCTTGC |
| *AT5G51890* | *AT5G51890* | Col-0 | *at5g51890-1* | SALK_086448.49.80.x | TAGATTCGACACGGTCAAACC | TTTACCTGAATCAAGCCCATG |
| | | Col-0 | *at5g51890-2* | SAIL_606_F08 | CGAAACCATTTCTTCTCTACCG | ATTCTTAGGAGACGAGCAGGC |
| *IRX8* | *AT5G54690* | Col-0 | *irx8-1* | SALK_014026.29.99.f | ATCATTACTCCGATCCCGAAG | TGAGTTGCGGGTTGATATCTC |
| | | | *irx8-2* | SALK_071351.39.55.x | CTGGAAAGGCAACAAGTCTTG | TCAACGGTGGAGAGAACAAAG |
| *PMR5* | *AT5G58600* | Col-0 | *pmr5-1* | SALK_034969.29.99.f | TCACGAAGAAGGTCAAAATGC | CTGACAATCGAACTCAGGCTC |
| | | | *pmr5-2* | SAIL_298_D05 | TATCAACGTGAGTTTCGACCC | GAGCCTGAGTTCGATTGTCAG |
| *ARR6* | *AT5G62920* | Col-0 | *arr6-2* | SALK_008866.55.00.x | TGTTGAGGAAAAATCAGTCGG | CTGCGAGTGAACAGGGTAGAC |
| | | | *arr6-3* | SALK_133123.21.20.x | TCTTCTGGGCCAAATCATATG | TACCGGGCATTGAGTAATCAG |

**Table S4: Oligonucleotides used for gene expression analysis.**

| Gene | AGI locus | Forward oligonucleotide | Reverse oligonucleotide |
|---|---|---|---|
| *PHI-1* | *AT1G35140* | TTGGTTTAGACGGGATGGTG | ACTCCAGTACAAGCCGATCC |
| *PR1* | *AT2G14610* | CGAAAGCTCAAGATAGCCCACA | TTCTGCGTAGCTCCGAGCATAG |
| *PR4* | *AT3G04720* | AGCTTCTTGCGGCAAGTGTTT | TGCTACATCCAAATCCAAGCCT |
| *PDF1-2* | *AT5G44420* | TTCTCTTTGCTGCTTTCGACG | GCATGCATTACTGTTTCCGCA |
| *LOX3* | *AT1G17420* | GCGGAGATTGTTGAAGCGTTT | GCCCCACACCTATTTCTACGGT |
| *PAD3* | *AT4G31500* | CAACAACTCCACTCTTGCTCCC | CGACCCATCGCATAAACGTT |
| *LTP3* | *AT5G59320* | GAAGAGCATTTCTGGTCTCAAC | GTTGCAGTTAGTGCTCATGGA |
| *PHI1* | *AT1G35140* | TTGGTTTAGACGGGATGGTG | ACTCCAGTACAAGCCGATCC |
| *WRKY33* | *AT2G38470* | ACGGCCAGAAAGTCGTTAAGG | CATGTCGTGTGATGCTCTCTCC |
| *CYP81F2* | *AT5G57220* | TATTGTCCGCATGGTCACAGG | CCACTGTTGTCATTGATGTCCG |
| *AT1G51890* | *AT1G51890* | CCAGTTTGTTCTGTAATACTCAGG | CTAGCCGACTTTGGGCTATC |
| *UBQ21* | *AT5G25760* | GCTCTTATCAAAGGACCTTCGG | CGAACTTGAGGAGGTTGCAAAG |
